# The mode and repeatability of thermal adaptation in gene expression in the seed beetle *Callosobruchus maculatus*

**DOI:** 10.64898/2026.07.29.741406

**Authors:** Alex F Hart, Alexandre Rêgo, Rike Stelkens, David Berger

**Author notes:** Corresponding author: David Berger Department of Ecology and Genetics Uppsala University, Uppsala 752 36, Sweden. These authors contributed equally (Alex F Hart, Alexandre Rêgo).

## Abstract

Transcriptional plasticity can maintain organismal homeostasis but its potential to buffer negative effects from climate change remains contentious and may rely on genetic adaptation. Theory predicts two general outcomes: genetic reinforcement, where adaptive plasticity is enhanced by genetic adaptation; or genetic compensation, where genetic adaptation reverses maladaptive plasticity. Here, we explored the prevalence, repeatability, and molecular basis of compensation and reinforcement of gene expression responses to temperature using experimental evolution in the beetle pest, *Callosobruchus maculatus*. We evolved lines from three genetic backgrounds under hot or cold conditions and compared them to ancestral lines after 80-135 generations. Cold adaptation was dominated by genetic compensation, revealing a highly repeatable mode of adaptation across backgrounds, however, the underlying genes and their biological functions were largely idiosyncratic. In contrast, heat adaptation showed a less consistent mode, combining compensation and reinforcement, but greater repeatability at the level of genes and their functions. Network analysis identified many heat reinforcement genes as hubs and strongly temperature dependent, suggestive of roles in orchestrated adaptive thermal plasticity, potentially explaining their high repeatability. Genomic analyses identified repeatable allele frequency changes at both hot and cold compensation and reinforcement genes during experimental evolution. However, these were not causally linked to changes in expression levels, suggesting trans-regulation. There was also no broader correspondence between the level of repeatability of sequence and expression evolution across gene categories. Our findings suggest that the predictability of evolution under climate change might critically depend on the thermal range and the level of the genotype-phenotype map that is studied.

## Introduction

Phenotypic plasticity is a potent mechanism by which organisms can respond to changes in their surroundings, but the speed and pervasiveness of climate change may render current plastic responses inadequate (1–5), necessitating rapid genetic adaptation (6–8). Yet, phenotypic plasticity is likely to play a central role in moderating genetic responses (9–14). Theory suggests that adaptive plasticity can act as a buffer for genetic adaptation by allowing organisms to survive initial bouts of environmental change (11,12,15–18). In this scenario, the genetic response to selection is expected to occur in the same direction as phenotypic plasticity, reinforcing the initial plastic change towards the new phenotypic optimum. From hereon we refer to this scenario as “*genetic reinforcement*”. Alternatively, if plasticity is maladaptive in the new environment, it could facilitate (or necessitate) rapid genetic adaptation by exposing poorly adapted alleles to selection (3,19–22). In this case, the genetic response is expected to occur in the opposite direction from the initial plastic response, reverting the phenotype back towards the ancestral state. From hereon we refer to this scenario as “*genetic compensation*”.

Both reinforcement and compensation entail genetic adaptation, but their causes and consequences differ. Adaptive plasticity, implicit in the reinforcement scenario, is likely to weaken selection on environmentally sensitive phenotypes by reducing environmental mismatch. Such plastic responses can maintain population size, leaving opportunity for subsequent genetic adaptation (12,15,23). However, by weakening selection, adaptive plasticity may under certain conditions slow genetic responses to environmental change and increase extinction risk (24–27). The compensation scenario is instead more likely to entail stronger selection at genes encoding maladaptive plasticity. Such strong selection can lead to rapid genetic adaptation given abundant standing variation (3,12,19), but also to population decline in the presence of genetic and demographic constraints (26,28,29). Understanding the extent to which plasticity is adaptive or maladaptive, and whether environmental adaptation proceeds via genetic reinforcement or compensation, can thus inform predictions of demographic and evolutionary responses to climate change (9–11,15,27,30–34).

Studies of climate change adaptation are increasingly employing transcriptomic data to explore the molecular basis of adaptation (30–32,35,36). Gene expression responses are particularly suitable for asking questions about genetic compensation and reinforcement as thousands of molecular phenotypes in the form of RNA expression levels can be interrogated across the entire transcriptome. Gene expression further constitutes the intermediary between fitness-related life-history phenotypes and variation at the DNA sequence level and can provide information on the effects of environmental perturbations on the genotype-phenotype map (37–44).

The growing consensus from recent transcriptome studies suggests that genetic compensation is the more common mode of adaptation, with the majority of ancestral plastic responses being reversed during longer term adaptation (22,45–49,49–56). However, comparisons of the few studies at hand also reveal disparities in transcriptional responses and the underlying reasons are not well understood. This may in part be a result of different organisms and stressors being assayed, preventing a systematic overview of the link between molecular plasticity and adaptation. In particular, while a pressing objective is to predict future responses to climate change across different taxa and geographic regions using omics data (6), the factors that govern the repeatability of adaptation in gene expression remain poorly understood (30–32). Moreover, parallel changes in the expression of genes across populations or experimental replicates could naively be taken as evidence for responses to selection at causal genes, but such changes can also simply be by- products of adaptation at a few central regulatory loci that affect the expression of multiple downstream genes across the transcriptome (52,57–59). Distinguishing between whether adaptation of transcriptome plasticity proceeds via allele frequency changes at a few trans-regulatory loci with large fitness effects, or at many loci of small effect located within or around the region of differentially expressed genes, is central to understanding adaptive potential and the repeatability of evolutionary trajectories (27,41,42,60–62).

To explore how natural selection operates to promote adaptive gene regulation in new climates, we leveraged long-term experimental evolution in the seed beetle *Callosobruchus maculatus* (Figure 1). We exposed replicated populations derived from three different geographic origins to cold or hot temperature for 80-135 generations while keeping the founding populations at the ancestral temperature. We subsequently assayed gene expression to quantify ancestral thermal plasticity and evolutionary responses, allowing us to assess the prevalence of genetic compensation and reinforcement during adaptation to cold and hot temperature. To explore if transcriptome evolution was shaped by different selection pressures and genetic architectures for reinforcement versus compensation, we used DNA pool-seq data to infer historical purifying selection and directional selection during experimental evolution in genomic regions within and upstream of differentially expressed genes. Our replicated experimental evolution design, including genetic backgrounds from three different geographic origins, enabled us to assess if historical contingency influenced the mode and repeatability of gene expression evolution at hot and cold temperature, and whether these responses were reflected in underlying coding sequence evolution and higher-level life-history phenotypes.

**Figure 1:**
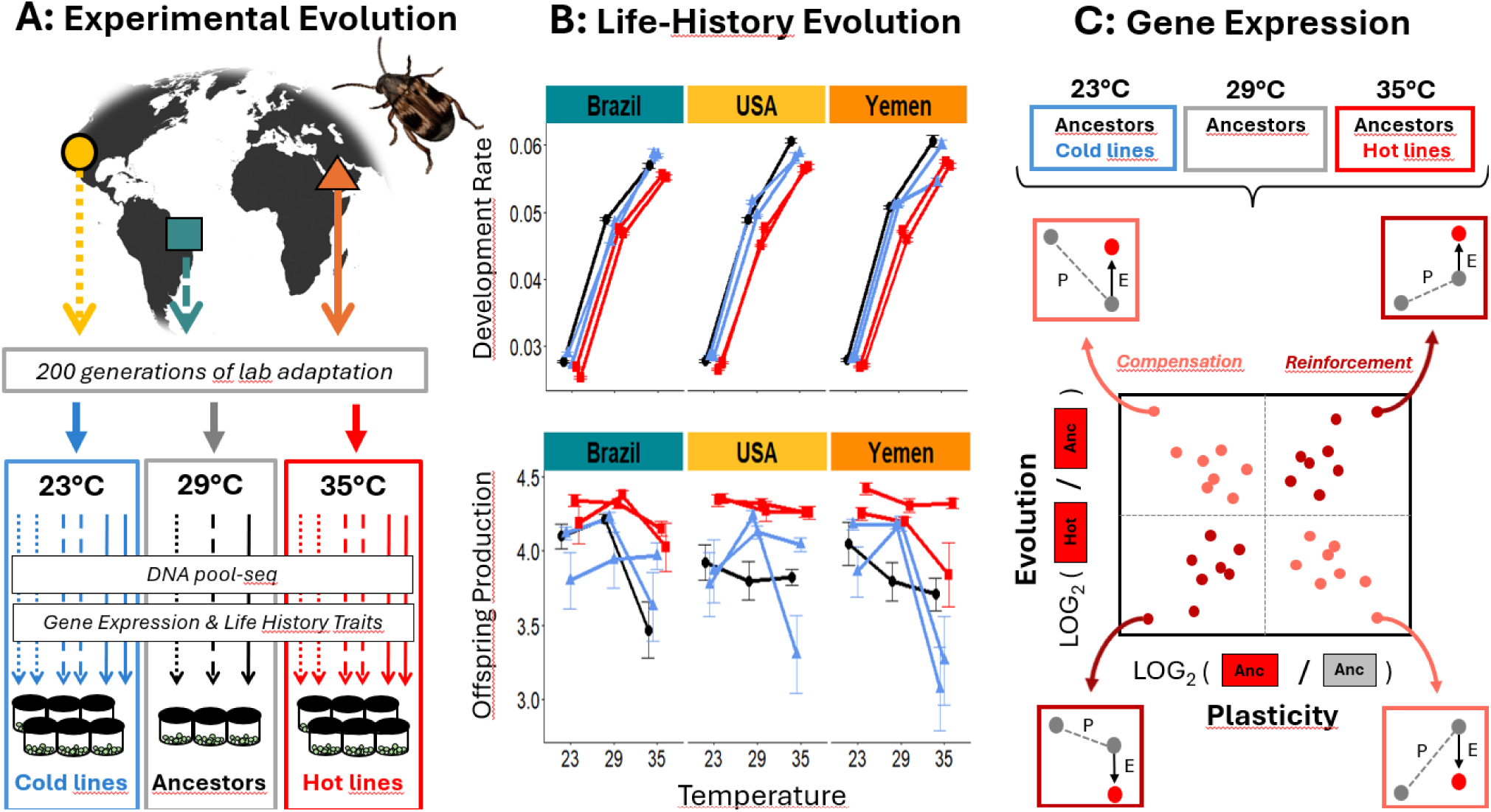
Study overview. A) Three genetic backgrounds (Brazil = teal, USA = yellow, Yemen = orange) were kept at lab conditions for ca. 200 generations after which replicates of these populations were exposed to experimental evolution at 23°C (cold lines) or 35°C (hot lines). Ancestral founders were maintained as single populations at the original benign 29°C. All lines were assayed for gene expression and life-history phenotypes after 80-135 generations in a common garden design. B) Life history data reanalyzed from von Schmalensee et al. (72) demonstrates that phenotypic evolution shows signs of genetic compensation (or “countergradient variation”): Cold temperature slows development, and cold lines show faster development than hot lines; hot temperature decreases (log_e_) offspring production, and hot lines show greater offspring production than cold lines (details in Supplementary 1). C) Gene expression data from pooled female abdomens was used to estimate thermal plasticity in ancestors reared at all three assay temperatures, and genetic evolution by comparing evolved lines to ancestors at cold and hot temperature. Responses of differentially expressed genes were classified as either genetic compensation (plastic and evolved response in opposite directions, light red color) or genetic reinforcement (plastic and evolved response in same direction, dark red color). In the example shown for heat adaptation. Compensation and reinforcement genes (individual points) were classified based on showing significant differential expression for both plastic and evolved changes. The identity, frequency, biological functions, and sequence variation of compensation and reinforcement genes were compared across genetic backgrounds to estimate the repeatability and mechanistic basis of cold and heat adaptation in gene expression.

## Materials and Methods

### Experimental evolution

With origins in central Africa, the seed beetle *Callosobruchus maculatus* has colonized tropical and subtropical regions throughout the world, where it is a widespread pest on fabaceous seeds (63–65). *C. maculatus* is well adapted to warm and dry climates and adults are facultatively aphagous, which allows them to infest seed storages where only larvae have access to food (63,66,67). Experimental evolution lines were established from three lab stocks originally sampled from Brazil, Yemen, and California (USA), respectively, that had been adapting to standard laboratory conditions (29°C, 55% RH) for ca. 200 generations prior to establishment of the lines. From each geographic origin (from hereon: “genetic background”), we established two replicate lines that were placed at 35°C (six hot lines), and two replicate lines placed at 23°C (six cold lines). We also maintained the original lab stock at the ancestral temperature of 29°C (three “ancestors”), for a total of 15 experimental populations (Figure 1A). Relative humidity was maintained at 50-55% throughout. These temperatures span the natural range of temperatures at each geographic site (68). Evolution lines were maintained by, at each generation, moving 600 newly emerged adults onto 250 ml of black-eyed beans (*Vigna unguiculata*) in 1 litre glass jars. The ancestors were maintained by moving 300- 400 newly emerged adults onto 150ml of black-eyed beans. All lines were transferred to fresh jars of beans at the peak of emergence in each generation, typically 3-5 days after the emergence of the first beetle, reducing directional selection on development time. Further details in: (69–71).

### Experimental design

To separate plastic from genetic changes in gene expression, all populations were first reared in a common garden for 2 generations at ancestral 29°C to remove parental (non-genetic) effects. Beans bearing newly hatched larvae (F0 generation) were transferred from each respective evolution regime to the common 29°C. Emerged F1 adults were then allowed to lay eggs for 48h on fresh beans, which were then split between 23, 29, and 35°C. Emerging virgin F2 adults were collected at the day of eclosion. For each evolution line and assay temperature, five males and five females were placed together in a 90mm petri dish and allowed to mate and reproduce, with access to 20 black-eyed beans as egg-laying substrate. Two such petri-dishes were set up per evolution line and assay temperature combination. For the three ancestors, each represented by only a single stock population, we set up four replicate petri-dishes. Adults were kept together for 23h at 29 and 35°C or 46h at 23°C; the longer duration at the cold temperature compensates for the thermodynamic slowdown of female reproduction and allows comparisons between females of similar biological age (70). Females were then flash-frozen in liquid nitrogen and dissected under sterile conditions, with abdomens retained for RNA extraction. Females from each pair of dishes were pooled into a single RNA library (10 females per library), yielding one library per evolution line and assay temperature combination. For ancestors, the four dishes were pooled into two libraries per ancestor and assay temperature combination. The full experiment was conducted in 2022 (after 80 generations of cold and 120 generations of hot experimental evolution) and repeated in 2023 (90 and 135 generations, respectively; Figure 1C). Thermal plasticity in gene expression was estimated from ancestors reared at all three assay temperatures. Cold and heat adaptation were estimated by comparing ancestors to cold lines at 23°C and to hot lines at 35°C, respectively. The full design yielded 60 RNA libraries.

Additionally, data on phenotypic adaptation in life-history traits from the siblings of the females chosen for gene expression were re-analyzed from von Schmalensee et al. (2026) (72) and are presented graphically in Figure 1B (Methods and Results in Supplementary tables 1A and 1B).

### RNA extraction and sequencing

RNA was extracted from each pooled sample using a Qiagen RNeasy Mini Kit with β-mercapto- ethanol added to the lysis buffer and an on-column DNase treatment with Qiagen RNase-free DNase kit. Tissue lysis was performed with a bead mill, using two stainless steel beads at 29 Hz for 90s. RNA was eluted in 2x 30-50µl water. Samples with insufficient RNA concentration underwent an additional clean-up step with the Qiagen RNeasy Mini Kit. RNA samples were sequenced through the SNP&SEQ Technology Platform at SciLife Lab Uppsala. Libraries were prepared using the TruSeq Stranded mRNA Library Preparation Kit with polyA selection and sequenced on a NovaSeq 6000 platform to generate paired-end 150 bp reads. Samples from each experimental year were sequenced on separate flow cells.

### Pre-processing of Sequencing Data

After data download, FastQC (v0.11.9) was used to visualise library quality statistics, as well as MultiQC (v1.1). To trim low confidence base calls and filter poor quality individual reads from libraries, Trimmomatic (v0.39) in paired-end mode was used with the following settings:

ILLUMINACLIP: TruSeq3**-**PE:2:30:10:2:true, LEADING:20, TRAILING:20, SLIDINGWINDOW:5:20, MINLEN:20. Read pairs which passed initial QC were aligned to the reference genome using HISAT2. The GCA_949361665.2 version of the *C. maculatus* reference genome was used (73). HISAT2 was run with default settings in *–very-sensitive* mode for increased mapping rates. To prepare the data for feature counting, SAMtools (v1.17) was used to sort each library, filter for unmapped reads, and indexing. The number of reads aligned to each feature was counted by HTSeq- count (v2.0.2). Union mode was used with default settings for paired-end data. Overall, this resulted in 9-35 million uniquely mapped reads per library.

### Differential Gene Expression Analysis

Read count data was processed using the R package edgeR v4.2.0 (74). Separately for each year, read counts were filtered to remove genes with low expression using the filterByExpr() function with a design of (∼0 + regime + origin + assay). Of 21,233 input genes, 12,961 passed this initial filtering step in 2022 and 12,230 in 2023, of which 11,923 were shared between years. To focus on genes with consistent expression across years, only genes with a Pearson correlation coefficient ≥ 0.25 between 2022 and 2023 log-CPM values were retained, yielding 8,219 genes. A further filter requiring CPM ≥ 10 in at least four samples left 6,739 genes for downstream analysis. Normalization factors were recalculated using calcNormFactors() after each filtering step.

To estimate ancestral plasticity, we used pairwise contrasts between the ancestral temperature (29°C) and each assay temperature (23°C or 35°C), fit separately for each of the three genetic backgrounds using a design of (∼0 + assay temperature + year). For a “global” estimate of ancestral plasticity pooling across genetic backgrounds, we fit the same pairwise contrasts using a design of (∼0 + assay temperature + genetic background + year), in which background was included as an additive fixed effect. To estimate genetic adaptation, we compared ancestors to cold lines at 23°C and ancestors to hot lines at 35°C, again fit both separately for each genetic background (∼0 + regime + year) and globally across backgrounds (∼0 + regime + genetic background + year). For each model, common, trended, and tagwise dispersion were estimated using estimateDisp(), reads were fit to generalised linear models using glmFit(), and differentially expressed genes were identified by likelihood ratio tests using glmLRT().

### Ancestral Thermal Plasticity: classification and repeatability

To understand ancestral plastic responses to temperature, the log fold-change and *p*-value for each gene were extracted for the calculation of cold plasticity (23 vs. 29°C) and heat plasticity (35 vs 29°C) for each genetic background separately, using only samples from ancestral populations. Significantly differentially expressed genes were classified into one of the following four categories: “Antagonistic” (showing opposing responses to hot and cold assay temperature), “Synergistic” (showing concordant responses to hot and cold assay temperature), “Heat-Limited” (only responding significantly to hot assay temperature), and “Cold-Limited” (only responding significantly to cold assay temperature). We then tested whether the overlap of the classified genes across the three genetic backgrounds was larger than expected by chance using Hypergeometric tests for multiple intersections in the package *SuperExactTest* (75). Note here that because we were mainly interested in analysing patterns of congruent and opposing responses in plastic and genetic responses, and not the significance of particular candidate genes, we did not correct p-values for multiple testing in any analyses, noting that false positives may have reduced the relative strength of observed overlaps.

### Thermal Adaptation: classification and repeatability

To understand genetic responses to temperature, the log fold-change and *p*-value for each gene were extracted for the calculation of cold adaptation (Cold-lines versus ancestors reared at 23°C) and heat adaptation (Hot-lines versus ancestors reared at 35°C) for each genetic background separately. Significantly differentially expressed genes were classified into the same four categories as done for the plastic responses (“Antagonistic”, “Synergistic”, “Heat-limited”, and “Cold-limited”) and overlap between genetic backgrounds was again tested.

### Genetic Compensation versus Genetic Reinforcement

To quantify the relationship between plastic and genetic responses to temperature on each genetic background, log fold-changes of genes that showed significant plastic and genetic responses to temperature were compared, for hot and cold adaptation separately. We classified genes into two major categories corresponding to the two modes that adaptation can take in relation to phenotypic plasticity; compensation, signified by the genetic response going in the opposite direction relative to the plastic response; and reinforcement, signified by genetic and plastic responses going in the same direction (Fig. 1C). We then quantified the mode of adaptation for each genetic background at hot and cold temperature by i) counting the number of genes that fell into each of the two categories, and ii) by calculating the correlation between the plastic and genetic responses (log-fold changes), with a negative correlation signifying genetic compensation and a positive correlation signifying genetic reinforcement. We then compared if the mode of gene expression adaptation was qualitatively similar across cold and hot temperature and across the three genetic backgrounds.

The correlation between plastic and genetic responses is biased by shared measurement error in the ancestral samples assayed at 23°C or 35°C, used to calculate both plasticity and adaptation. For our calculations of log fold-changes, this bias makes the correlation downwardly biased *a priori*. However, the strength of this bias is proportional to the magnitude of the measurement error relative to the true effect, and therefore becomes marginal when looking only at genes with statistically significant responses (76). Nevertheless, to remedy this issue further and verify the results from the global analyses, we leveraged our replicated experimental design to calculate independent and unbiased estimates of the correlation between plastic and genetic response vectors for each evolution line in each experimental year (a total of 12 estimated correlations each for heat and cold adaptation). In brief, to generate these estimates, one technical replicate of the ancestor at the hot temperature was used to calculate the plastic response (e.g. ancestor at 35 vs 29°C) while the other technical replicate from the same year was used to calculate the genetic response vector (e.g. heat-evolved line versus ancestor at 35°C) such that the two estimated response vectors did not share measurement error. We then rotated the ancestral samples in such a way that all samples were used once to calculate a plastic response vector and once to calculate a genetic response vector, which were used in different calculations of the correlation (see schematic in Supplementary Table 2). For each genetic background, we based the calculations only on genes that showed significant plastic and genetic responses in the background-specific analyses.

### Gene Ontology

Gene Ontology (GO) analysis was performed for each of the four classes of genes (Cold/Heat : compensation/reinforcement) separately, using the R package clusterProfiler (77). The GO term universe was adapted from the *C. maculatus* set generated by Sayadi et al. 2016 (78) and enrichment was assessed using hypergeometric tests. We tested whether GO terms with a minimum of two gene hits showed overlap between all three genetic backgrounds exceeding expectation using a permutation-based approach. Genes were resampled from the annotated gene universe while preserving gene list sizes. GO enrichment was then recomputed for each resample, and overlap among the N number of top-ranked GO terms was calculated (N = 10/20/40, ≥2 genes per term). We then calculated a test statistic defined as the number of shared terms divided by the number of terms on the background with fewest terms (setting the upper limit for the overlap). This fraction thus ranges between 0 and 1, irrespective of the length of gene lists. The empirical p-value was defined as the proportion of permutations producing overlap equal to or greater than the observed overlap.

### Modularity and co-regulation of compensation and reinforcement genes

To better understand the underlying genetic architecture of compensation and reinforcement genes, we performed a gene co-expression network analysis using the WGCNA package for R (79). Networks were built using only the 36 ancestral sampled assayed across the three temperatures (23, 29, and 35°C), so that module structure reflected the ancestral transcriptomic response to temperature independent of any evolved changes. After confirming that all samples and genes passed WGCNA’s quality checks, 6,739 genes were retained. A signed adjacency matrix was constructed using a soft-thresholding power of 9, chosen from the scale-free topology fit criterion (signed R² ≥ 0.85). The adjacency matrix was transformed into a topological overlap matrix (TOM), and the corresponding dissimilarity (1 – TOM) was used as input for average-linkage hierarchical clustering. Modules were identified from the resulting dendrogram with the dynamic tree-cut algorithm (deepSplit = 2, minimum module size = 30 genes), and modules with eigengene correlations above 0.75 were subsequently merged (eigengene-dissimilarity cut height = 0.25), yielding the final set of co-expression modules.

To identify modules associated with thermal plasticity, the eigengene of each merged module was correlated with assay temperature (Pearson’s correlation, with significance assessed using Student’s t-distribution). Modules with a significant temperature–eigengene correlation were retained as temperature-responsive and were the focus of subsequent analyses. We then tested whether hot and cold compensation and reinforcement genes were over- or under-represented within these modules relative to the 6,379 genes retained in the network.

To assess whether genes in each of the four categories tended to occupy peripheral or central (hub) positions within the network, we quantified each gene’s intramodular connectivity as its signed module membership (kME) to its own assigned module eigengene, defined as the Pearson correlation between the gene’s expression and that module’s eigengene; higher kME indicates a more central, hub-like position.

To test whether compensation and reinforcement genes lie within locally co-regulated genomic domains, we asked how strongly each gene’s expression correlated with that of its neighbours as a function of the genomic distance between them. Using the ancestral-sample co-expression network (see above), we computed the absolute correlation (|r|) for every same-scaffold gene pair and binned pairs by the distance between their midpoints. A correlation that decays with distance is expected given local co-regulation. Comparing the correlation and its decay computed across all gene pairs to that computed between pairs belonging to the same gene category (e.g. heat compensation) then tests whether local co-regulation is specific to genes of the same category rather than a generic local genomic domain. We compared each category’s observed correlation decay to a size-matched null sample of 1,000 equally sized random gene sets, taking the 2.5th and 97.5th percentiles per bin as a 95% envelope. Additionally, as a distance-independent reference, we calculated the mean |r| of 200,000 randomly drawn pairs of genes situated on different scaffolds (the cross-scaffold background).

### Estimates of purifying selection on compensation and reinforcement genes

We estimated the strength of purifying selection on the underlying DNA sequences of compensation and reinforcement genes using previously published DNA pool-seq data (69). This data was taken at generations 57-62 from the 12 evolved lines, and–as a surrogate estimate of the ancestral populations–from the 6 heat-evolved lines sampled at generation 3. We assessed the strength of purifying selection by calculating the ratio between the frequency of non-synonymous to total genetic polymorphism, π_N_/(π_N_+π_S_), for each gene, expecting a higher ratio for genes which expression is under weaker stabilizing selection. We then compared the frequency distributions of this estimate for compensation and reinforcement genes for heat and cold adaptation. We tested for differences in π_N_/(π_N_+π_S_) ratio between reinforcement and compensation genes using non-parametric paired Wilcoxon rank-sum tests with the two median ratios estimated within a given pool-seq sample as paired observations. In line with theoretical expectations (80), pool-seq samples from ancestral and evolved lines did not show any obvious differences in π_N_/(π_N_+π_S_) (Supplementary Figure 14). We therefore performed the analysis based on all 18 pool-seq samples as the level of replication. The test was performed for genes identified for hot and cold adaptation separately.

### Estimates of directional selection on compensation and reinforcement genes

We used allele frequency change between ancestral and evolved populations to assess directional selection during experimental evolution. We explored: (i) whether allele frequency change at and upstream of compensation and reinforcement genes was repeatable across independent evolutionary replicates within and between genetic backgrounds, and (ii) whether the magnitude of allele frequency change was associated with changes in gene expression (i.e., log-fold expression change between ancestors and evolved lines), as expected if cis-regulation was prominent.

For each SNP in each population, we computed both the raw allele frequency change (***Δ***AF =*p*_1_ − *P_0_*) and the standardized allele frequency change 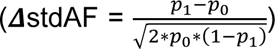, where p₀ and p₁ are the alternate-allele frequencies in the ancestral and evolved pools, respectively. The denominator is proportional to the expected variance in allele frequency change under drift or selection, so that *Δ*stdAF can be compared across SNPs of differing starting frequency within a population. Raw and standardized allele frequency changes were calculated both within the gene body and within the 1 kb region immediately upstream of the transcription start site as a likely target for cis-regulation.

To quantify repeatability of allele frequency change between two replicate populations of the same thermal regime, we used raw allele frequency change. To ensure that repeatability estimates reflected shared standing variation rather than fixed differences, we restricted these analyses to SNPs that were polymorphic in the ancestors of both compared populations. Repeatability was then quantified in two ways. First, we computed the Pearson correlation between *Δ*AF vectors of the two replicate populations. Second, we estimated the observed and expected overlap of outlier SNPs detected independently in the two populations. Outliers were identified as those whose allele frequency change exceeded the change expected by drift (p<0.05), parameterized by previously obtained estimates of *Ne* in these populations (69).

Finally, we tested if the magnitude of allele frequency change at a gene was associated with its position in the co-expression network (kME) or with its evolved log-fold expression change using linear regression applied separately for each genetic background.

## Results

### Ancestral Thermal Plasticity in Gene Expression

Across all three ancestral backgrounds, 3491 loci (out of 6739 analyzed) were differentially expressed in response to heat, with a significant enrichment of genes related to protein synthesis and heat shock. For the cold response, 1605 loci were differentially expressed, with an enrichment of genes related to sugar and nucleic acid metabolism. Analyzing each genetic background separately showed that observed overlaps in expression responses between backgrounds were greater than expected by chance for all classes of genes, but the odds-ratio was higher for Antagonistic and Heat-Limited genes compared to Synergistic and Cold-Limited genes. Heat-limited genes were also the most abundant category making up 50% or more of all significant genes on each background (Supplementary Figure 1, Supplementary Table 3).

### Thermal Adaptation in Gene Expression

The global analysis across all three backgrounds found evidence for heat adaptation at 1660 loci, with ontologies such as cell adhesion, nucleic acid biosynthesis, glycoproteins, cytoskeletal structure, protein synthesis, and sugar metabolism. The corresponding analysis for cold adaptation found 792 significant loci, showing enrichment for sugar and carbohydrate metabolism, protein degradation, cytoskeletal structure, and nucleic acid metabolism. Heat- and Cold-limited genes generally showed significant overlap between backgrounds, but the overlap was greater for genes involved in heat adaptation (Supplementary Figure 2). There were few Antagonistic and Synergistic loci detected, noting that this result is in part due to the more stringent selection criteria of these genes (heat- and cold-limited genes require only a single significant test to appear on a background, while antagonistic and synergistic genes require two). Nevertheless, the fact that there were more Synergistic than Antagonistic genes found on each of the three backgrounds suggests a modest role for antagonistic pleiotropy in the observed adaptation to hot and cold temperature (Supplementary Figure 2, Supplementary Table 4), as previously observed at the level of SNPs in these lines (69).

### Genetic Compensation versus Genetic Reinforcement

Genes showing a plastic response were more likely to also show a genetic response to temperature in both the global analysis (Figure 2A) and on each background when analysed separately, with the exception of Brazil lines adapting to heat (Supplementary Figure 3). In the global analysis, we detected 280 loci with both significant plastic and genetic responses to cold, with 230 (82%) of these responses classified as genetic compensation, showing plastic and genetic responses in opposing directions (correlation between log-fold changes = -0.64, p < 0.001). This suggests that maladaptive transcriptional plasticity plays a relatively large role in cold adaptation. Interestingly, for heat adaptation, almost half of the genes (372/804: 46%) showed evidence of genetic reinforcement (correlation between log-fold changes = -0.39, p < 0.001), suggesting that the mode of transcriptome evolution differs at hot and cold temperature (Figure 2B).

**Figure 2:**
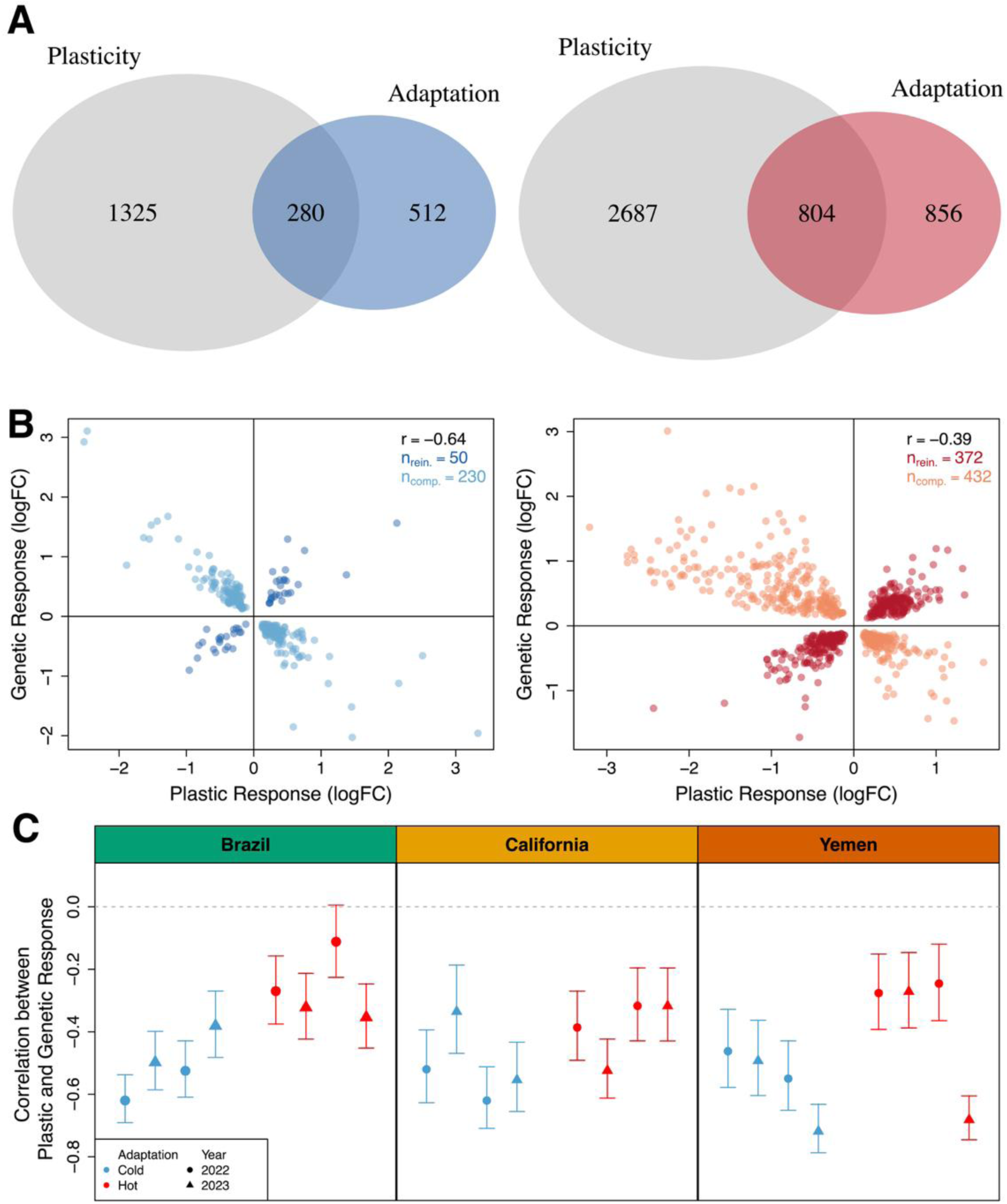
Genetic Compensation vs. Genetic Reinforcement during heat and cold adaptation. A: There was significant overlap between plastic (grey) and genetic responses to both cold (blue) and heat (red) in the global analysis. B: 82% (230/280) of the genes responding both genetically and plastically to cold in the global analysis showed signs of genetic compensation, whereas compensation and reinforcement where about equally likely for the 804 loci involved in heat adaptation C: The 24 independently estimated correlations between plastic and genetic response vectors based on each of the 12 evolved lineages’ responses measured in 2022 and 2023. Negative correlations designate compensation and positive correlations reinforcement as the main mode of adaptation (see panel B). Genetic compensation is a common and highly repeatable mode of cold adaptation in gene expression, whereas the mode of heat adaptation encompasses a mix of compensatory and reinforcing responses. Means and 95% CIs are based on correlations between genetic and plastic response vectors for each sample estimate, including only genes with both genetic and plastic responses being statistically significant in the background-specific analyses.

To verify these patterns, we first estimated the correlation between plastic and genetic log-fold changes independently for each population and year. We then performed a nested ANOVA with these 24 correlations as the response variable and population replicates as paired observations (six pairwise comparisons across evolution regimes) to test for differences in the mode of heat and cold adaptation. This showed that, while genetic compensation (negative correlation) was the overall more common mode of temperature adaptation, its prevalence differed between hot and cold lines (F_1,6_ = 10.4, P = 0.018, Figure 2C), with genetic compensation being more pronounced during cold adaptation. Similarly, the fraction of compensation genes relative to reinforcement genes was greater for cold adaptation on all three genetic backgrounds when analyzed separately (Brazil: χ^2^= 288.5, p < 0.001; USA: χ^2^= 48.4, p < 0.001; Yemen: χ^2^= 52.3, p < 0.001, Figure 3).

**Figure. 3.**
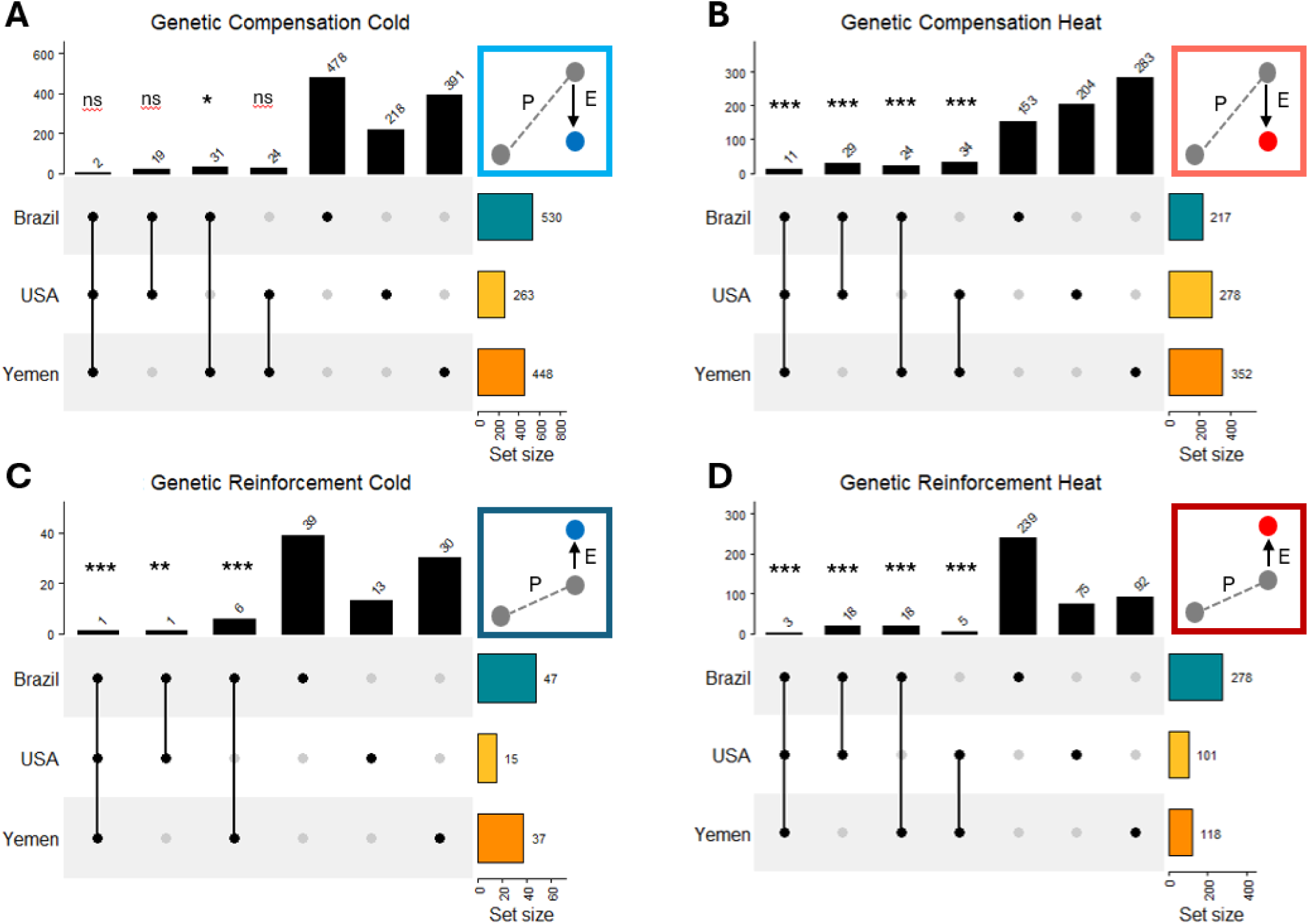
Overlap across genetic backgrounds of compensation and reinforcement genes. Upset plots showing gene re-use when adaptation proceeds via compensation (A, B; Plastic and Evolved response in opposite directions) or reinforcement (C, D; Plastic and Evolved response in the same direction). At hot temperature (B, D), both modes of adaptation show greater overlap than expected by chance. At cold temperature (A, C), however, only reinforcement shows greater overlap, whereas backgrounds do not share more genes involved in cold compensation than expected by chance. Hence, while the pattern of cold compensation was the most pronounced and repeatable mode of adaptation with many differentially expressed genes on all three backgrounds, the specific genes involved tend to be unique for each background. Asterisks indicate level of significance for overlaps (n.s. p > 0.05, * p < 0.05, ** p < 0.01. *** p < 0.001).

### Repeatability of the Genes underlying Compensation and Reinforcement

While genetic compensation was a highly repeatable mode of molecular adaptation to cold, very few of the involved genes were shared between genetic backgrounds, with gene overlaps generally not greater than expected by chance. In contrast, genes involved in cold reinforcement and heat adaptation showed much greater overlaps than expected by chance (Figure 3A-D, Supplementary Figure 4, Supplementary Table 5).

Despite 263-530 cold compensation genes detected on each background, only two were shared among all three backgrounds (Fig. 3A). One was annotated for Lysozyme 1, presumably linked to immune function, and another for Fatty acyl-CoA reductase, linked to cuticular hydrocarbons and water-proofing the cuticle, which is in line with the observed evolved increase in water loss of cold-adapted relative to ancestral and heat-adapted lines (70). A single out of the 15-47 cold reinforcement genes found on each background was shared among all three (Fig. 3C); Fumarylacetate hydrolase domain-containing protein 2 is overrepresented in mitochondria and plays a role in metabolism and catalytic activity, which is likely under strong selection at cold temperatures due to thermodynamic constraints on biological rates (81).

There were 11 heat compensation genes (out of 217-352) shared among backgrounds (Figure 3B). These showed annotations for functions such as larval development and digestion (Cathepsin L-like proteinase) and the oxidative stress response (Aldehyde dehydrogenase dimeric NADP preferring), which is likely under strong selection at hot temperatures (81). Interestingly, one of the 11 shared genes was Fatty acyl-CoA reductase, which was one out of the two cold compensation genes shared among backgrounds and linked to water-proofing the cuticle, which shows phenotypic divergence between heat- and cold-adapted lines. This implies potential antagonistic pleiotropy at this gene at hot and cold temperature.

There were three heat reinforcement genes (out of 101-278) shared among backgrounds (Figure 3D). These were annotated for; Carboxypeptidase E, involved in digestion, moulting and reproduction; PR domain zinc finger protein 16, which is a transcription factor with unknown and potentially very broad function in insects; and Ejaculatory bulb-specific protein 3 linked to male fertilization success in *Drosophila*.

### Repeatability of Biological Functions underlying Compensation and Reinforcement

The overlap in gene ontology was very strong for heat adaptation (Obs fraction/Exp fraction, Compensation: 0.40/0.01, P_perm_ < 0.001; Reinforcement: 0.21/0.03, P_perm_ = 0.002). Biological processes shared among all three backgrounds were related to cuticle and cell membrane function and catalytic activity for compensation, and to protein homeostasis and catalytic activity for reinforcement. Overlap was also seemingly strong for cold reinforcement (1.00/0.28, P_perm_ = 0.10), although the overlap was not greater than expectation due to the low number of reinforcement genes and GO terms entering the analysis (Figure 4); only two GO terms, catalytic activity and metabolic process, were found on the USA background, and these were also found for Yemen and Brazil. Interestingly, while cold compensation was associated with many GO terms on each background, only one GO term (heme binding) was shared among all backgrounds, and overlap was not greater than expectation (Obs/Exp: 0.05/0.01, P_perm_ = 0.17). This result recapitulates the low repeatability at the gene level and suggests that cold adaptation is dominated by finetuning of misexpression due to randomly accumulated conditionally deleterious alleles with effects that become exposed at cold temperature. In further support of this hypothesis, increasing the cut-off for the number of top-ranked terms on each background entering the analysis from 20 to 40, resulted in more overlap than expected also for cold compensation (Obs/Exp: 0.18/0.05, P_perm_ = 0.007, Supplementary Figure 5). Thus, while there is some functional overlap underlying cold compensation at the molecular level, the relative importance of these processes seem to differ between the backgrounds.

**Figure 4:**
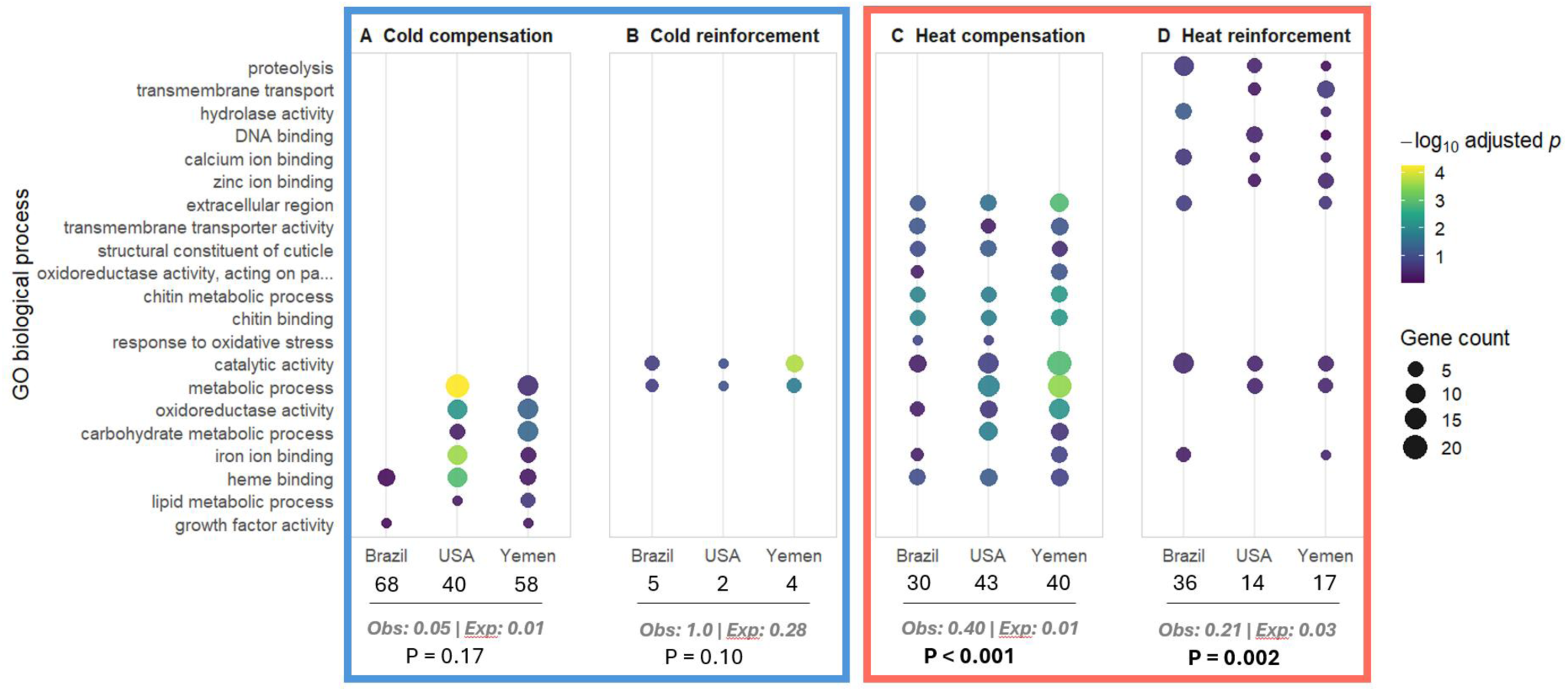
Overlap of gene ontology between genetic backgrounds. Shown are the top 20 biological processes with overlap between at least two genetic backgrounds and with at least two gene hits. The total number of GO terms for each background are shown on the x-axis. P-values refer to comparisons against simulated null-distributions. Observed and expected fractions of GO-terms that overlap all three backgrounds (relative to the number of GO-terms on the background with least annotated terms when below 20) are shown in grey. Heat adaptation resulted in more parallel responses in gene ontology compared to cold adaptation. In particular, cold compensation showed low repeatability across backgrounds while being associated with many GO terms within each background.

### Modularity and co-expression of compensation and reinforcement genes

We identified three large, temperature-associated gene modules; ’purple’, ’brown’, and ’green’ (Supplementary Figure 6). Gene Ontology over-representation tests recovered distinct functional programs for the three modules: the positively temperature-correlated purple module was enriched for DNA replication and nucleic acid/ATP binding, whereas the two negatively correlated modules (brown and green) were enriched for transport, protein-turnover, and cuticular functions (Supplementary Figure 7). Within each module, a gene’s module membership score (kME) was correlated with how strongly its expression depended on assay temperature (Supplementary Figure 6A). A higher kME also indicates a more central, hub-like gene; heat reinforcement genes tended to be more hub-like than the other three categories (median kME to their assigned module: cold compensation = 0.69, cold reinforcement = 0.64, heat compensation = 0.76, heat reinforcement = 0.85), a pattern that was most pronounced for reinforcement genes in the brown module, for which they were also enriched (Supplementary Figure 6B).

Beyond their positions within co-expression modules, we asked whether compensation and reinforcement genes are organized into locally co-regulated genomic regions. Across all four categories, a focal gene’s expression was most strongly correlated with that of its immediate genomic neighbours, and this correlation decayed with distance, confirming the expected local structure of the transcriptome (Figure 5, Supplementary Figure 8). The four gene categories differed, however, in how far local co-regulation extended and whether it exceeded random expectations. Heat reinforcement genes retained elevated co-expression out to intermediate and large distances (∼50kb to 5Mb; Figure 5) and exceeded their null expectations across this range (Supplementary Figure 8), suggesting strong co-regulation independent of physical genomic distance. Pairwise correlations were also stronger when computed only between genes within the heat reinforcement category (Figure 5). Heat compensation genes showed a similar but weaker pattern whereas cold adaptation genes showed co-expression at or below the null expectation beyond ∼50kb, consistent with these genes being dispersed across the genome and having more independent functions rather than being physically or functionally clustered. There was also no evidence that the elevated co-regulation of heat reinforcement genes (and to some extent heat compensation genes) could be explained by closer genomic distances, as genes within all four categories showed similar average distance to neighbours both within and outside their own category (Supplementary Figure 9).

**Figure 5:**
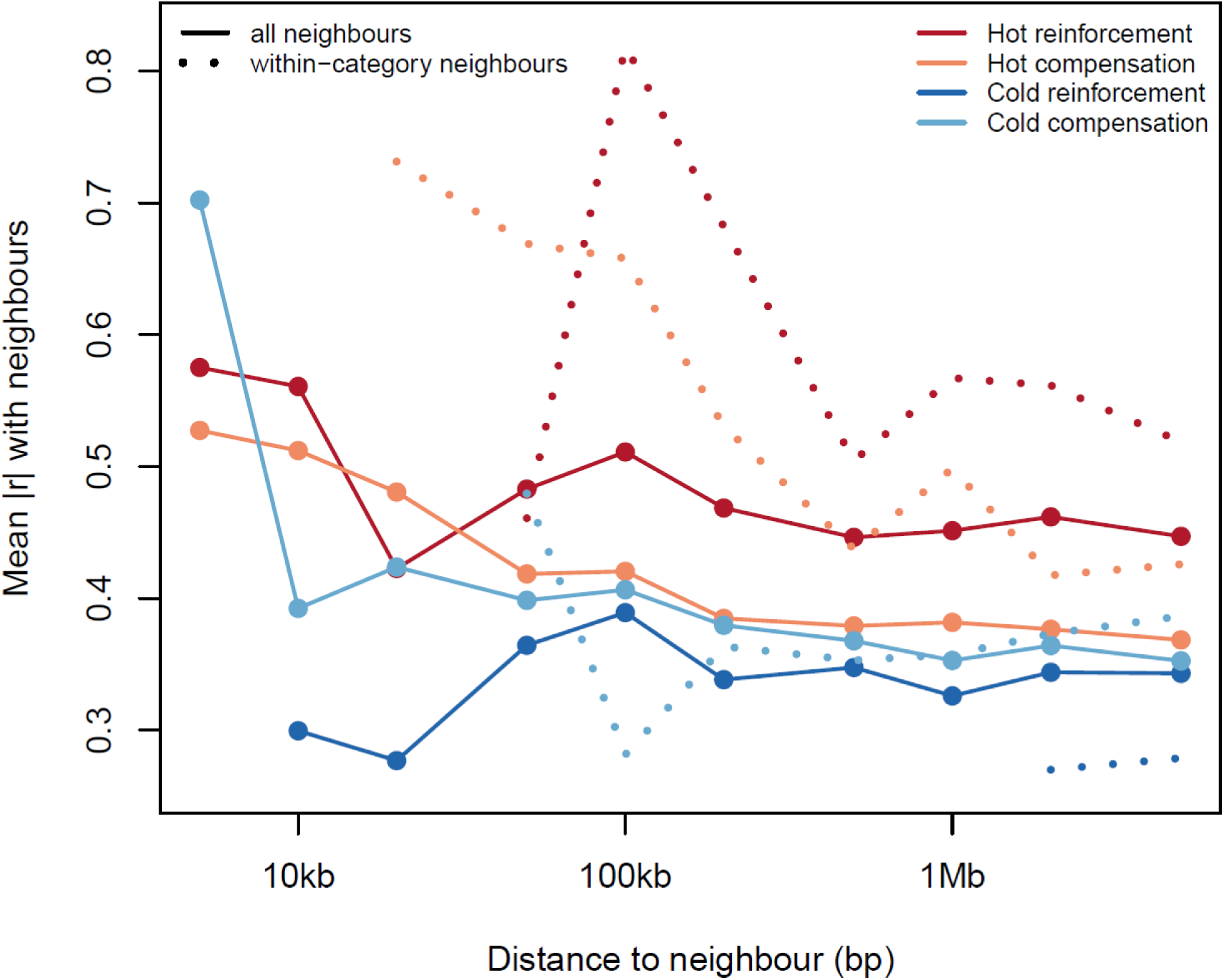
Co-expression of compensation and reinforcement genes. Mean absolute correlation of expression (|r|) between a focal gene belonging to one of the four categories and its same-scaffold neighbours, as a function of genomic distance. Solid lines, all neighbours; dotted lines, within-category neighbours. Correlations are computed from ancestral expression profiles. Both heat compensation and heat reinforcement genes show stronger pairwise correlations with genes within their own category, while genes involved in cold adaptation do not. Heat reinforcement genes show the overall strongest correlations and exceed the null expectation from random gene sets whereas correlations for genes belonging to the other categories fall within this distribution (see also Supplementary Figure 8).

These results are consistent with the hypothesis that some heat reinforcement genes occupy influential positions in the co-expression network and may help govern key molecular processes underlying orchestrated adaptive plasticity to heat regulated by trans-acting effects.

### Estimates of purifying selection on compensation and reinforcement genes

We compared the strength of purifying selection on reinforcement genes versus compensation genes (classified by the global analysis of differential expression across all three genetic backgrounds) using DNA pool-seq data (Figure 6, Supplementary Figure 10). We calculated the median value for π_N_/(π_N_+π_S_) for the two categories, for cold and heat adaptation genes separately, in each of the 18 pool-seq samples and then tested for differences using a simple paired t-test. For cold-adaptation, the median was greater for reinforcement than for compensation (t = 13.69, df = 17, p < 0.001, Δπ_N_/(π_N_+π_S_) = 0.040), indicating stronger purifying selection on compensation genes. For heat- adaptation, however, the median π_N_/(π_N_+π_S_) was slightly lower for genes involved in genetic reinforcement (t = 8.58, df = 17, p < 0.001, Δπ_N_/(π_N_+π_S_) = -0.011). These differences seem unrelated to the mean expression level of each gene class (F₁,₁₀₈₀ = 0.25, p = 0.62; Figure 6A). Overall genetic diversity (π_N_+π_S_) did not differ between compensation and reinforcement genes for cold adaptation (paired t-test, t = 0.03, df = 17, p = 0.98) but was lower at reinforcement genes for heat adaptation (t = 10.3, df = 17, p < 0.001; Supplementary Figure 11). Comparing ancestral and evolved samples, there was no sign that the π_N_/(π_N_+π_S_) ratios in genes underlying heat or cold adaptation evolved in any strong systematic fashion during the course of experimental evolution (Supplementary Figures 12-14), suggesting that adaptation in gene expression has been driven by changes at regulatory loci rather than by sequence evolution at the gene targets themselves.

**Figure 6:**
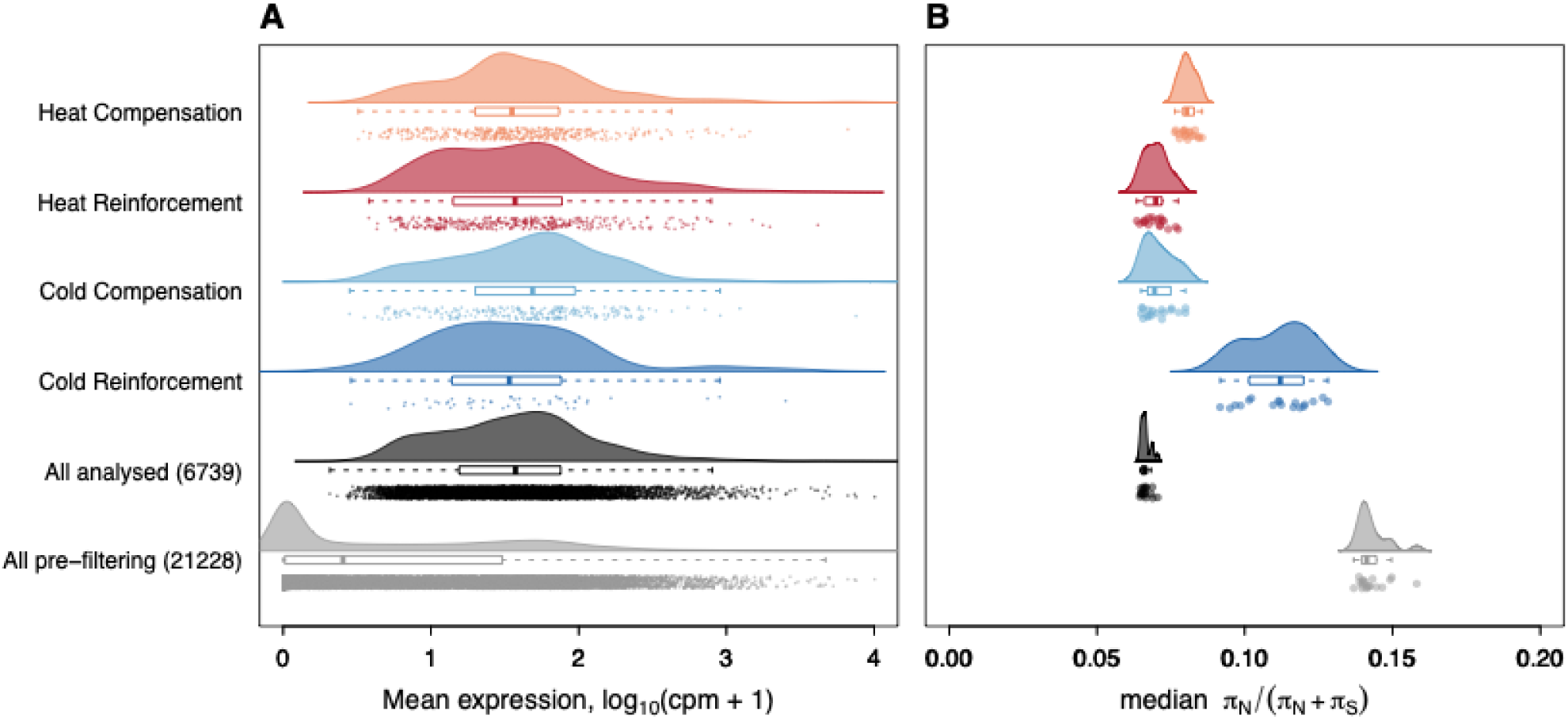
Purifying selection on compensation and reinforcement genes. A: The four classes of genes (light red: 432 Heat Compensation genes; dark red: 372 Heat Reinforcement genes; light blue: 230 Cold compensation genes; dark blue: 50 Cold Reinforcement genes) did not show any clear differences in expression levels compared to all the 6739 genes analyzed (black), but higher expression than all the 21233 genes pre-filtering (grey). B: Density plots based on median πN/(πN+πS) values for each of the 18 sequenced experimental samples. Heat reinforcement and cold compensation genes showed lower values of πN/(πN+πS) than heat compensation and cold reinforcement genes, indicating that the former have been under historically stronger purifying selection compared to other two gene categories. None of the four gene categories showed lower values of πN/(πN+πS) than the median of all 6739 analyzed genes, as could be expected for plastic genes with conditional expression (see also Supplementary Figures 10-12).

### Directional selection on compensation and reinforcement genes

To inspect this further, we looked for signals of directional selection on reinforcement and compensation genes using the pool-seq data. Standardized allele frequency changes within and upstream of compensation and reinforcement genes were generally of the same magnitude or smaller than estimates based on all analyzed genes (Supplementary Figure 15). Nevertheless, there was still evidence for significant directional selection within and upstream of several reinforcement and compensation genes, with many more outlier SNPs shared between replicate populations than expected by chance (Supplementary Figures 16-19). The overlap was stronger between replicates from the same genetic background, indicating a role for historical contingency in dictating sequence evolution (69). However, the strength of overlap between backgrounds was not greater for genes showing parallel expression changes compared to genes showing expression changes on only one of the two backgrounds (Figure 7A). Moreover, the strength of overlap in SNPs (strongest between backgrounds for cold compensation) was not related to the strength of overlap in differentially expressed genes or their functions (weakest for cold compensation genes).

**Figure 7:**
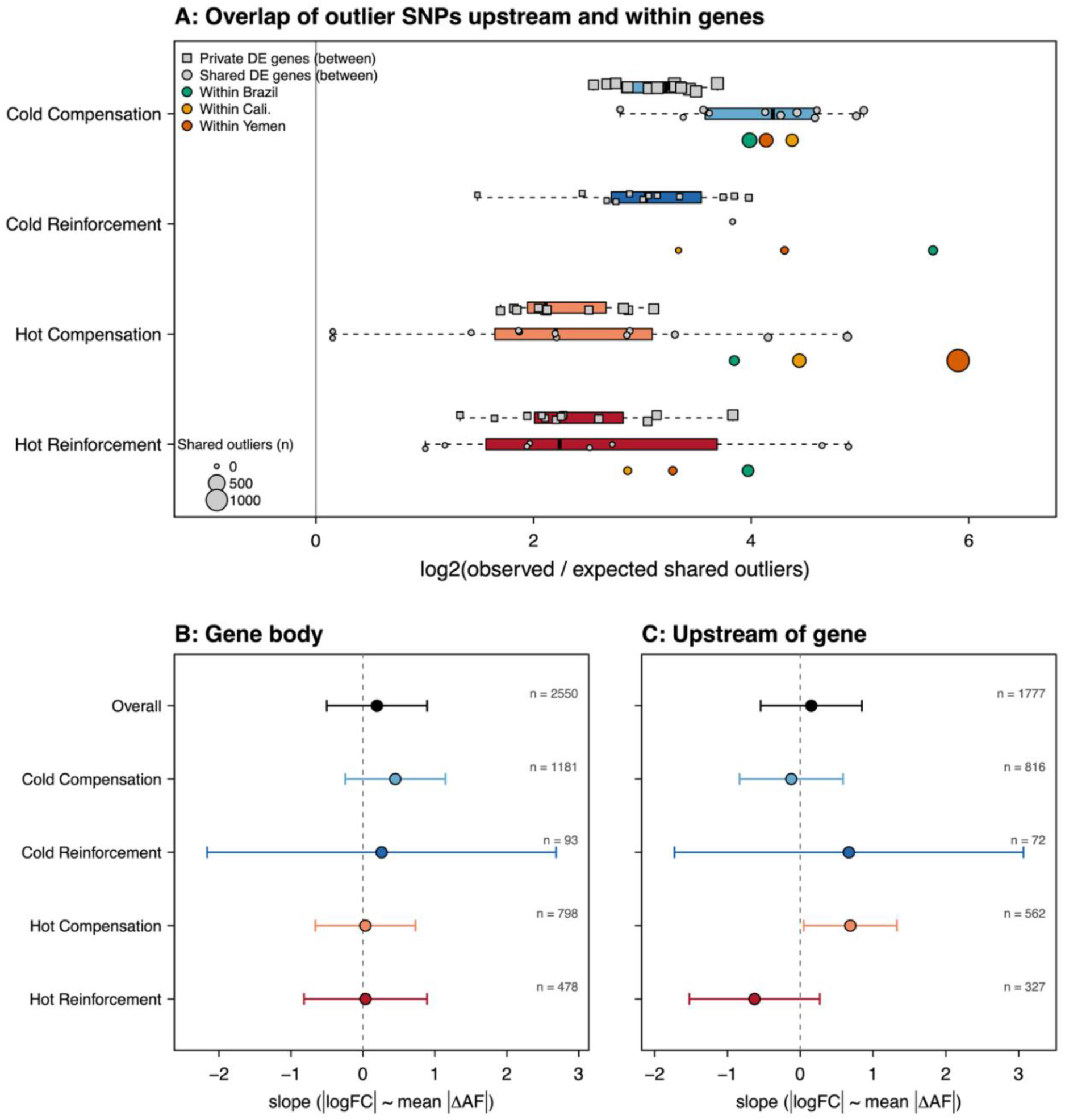
Directional selection on DNA sequences upstream and within compensation and reinforcement genes is repeatable but does not explain expression divergence. (A) Enrichment of shared outlier SNPs lying within the gene body and 1 kb upstream of the genes in each adaptation category. Overlap calculated for all pairwise population comparisons within each evolution regime and reported as log₂(observed number / expected number under independence); values of log₂ > 0 indicate more shared outlier SNPs than expected by chance. Box plots summarise the distribution across population pairs from different genetic backgrounds, shown separately for genes differentially expressed on a single background (private DE genes: grey squares) and those differentially expressed on both of the backgrounds being compared (shared DE genes: grey circles). Boxes give the median and interquartile range and whiskers the range. Colored circles show the overlap between the two replicate populations of the same genetic background. There is more overlap than epxcted by chance for all four gene categories. The overlap is greatest for cold compensation genes, which showed the weakest overlap for expression. Panels B and C show mean and 95% CI’s for regression slopes of the relationship between a gene’s allele-frequency change and the magnitude of its expression divergence during experimental evolution. Estimates are taken from the mixed model |logFC| ∼ mean|ΔAF| × category + background + (1 | gene), fitted separately for SNPs in the gene body (B) and 1 kb upstream (C). A slope indistinguishable from zero is the expectation under primarily trans regulation, whereas a positive slope may indicate a cis-regulatory contribution. n is the number of SNPs contributing to each estimate; “Overall” pools all four gene categories. There was no evidence for cis-regulation except for a weak, yet significant, positive slope for the analysis of SNPs upstream of heat compensation genes.

We calculated regression coefficients from the relationship between absolute log-fold change in expression on raw allele frequency changes upstream and within gene bodies (Figure 7B,C), expecting these to be positive if cis-regulation was a driver of gene expression levels. However, all coefficients were weak and non-significant except for the analysis of upstream heat compensation SNPs, which showed a weakly positive relationship with changes in gene expression. Thus, sequence evolution was at best only weakly related to the detected changes in expression.

## Discussion

Gene expression has been studied extensively in recent years to understand the molecular basis for how organisms respond to changing environments, with a growing body of work showing that evolutionary adaptation often reverses, rather than reinforces, ancestral plastic responses (45). This general pattern of genetic compensation suggests that ancestral plasticity is often maladaptive in new environments and that many populations must rely on genetic adaptation to achieve molecular homeostasis when faced with altered conditions. Our results generally agree with this picture as we observe pervasive genetic compensation across both hot and cold environments. However, we also show that the prevalence of genetic compensation versus reinforcement, and the involved genes and their repeatability, differ fundamentally between cold and heat adaptation. This suggests that our ability to predict evolutionary responses to climate may depend on at what level of biological organization we are asking our questions. In particular, we find that genetic compensation is a highly repeatable and dominating mode of cold adaptation across the transcriptome but entails different sets of genes across populations. Heat adaptation instead seems to entail a significant amount of genetic reinforcement, where genes with initially adaptive, but incomplete, plastic responses are repeatedly recruited across geographically separated populations.

### The mode and repeatability of thermal adaptation in gene expression

Cold adaptation was dominated by strong and highly repeatable pattern of genetic compensation: across all populations, most genes showing both plastic and evolutionary responses exhibited opposing changes in expression, indicating that ancestral plasticity under cold stress is often maladaptive and subsequently corrected through genetic evolution towards ancestral state. This result is also in line with our analyses of DNA pool-seq data, suggesting that cold compensation genes have been under the strong purifying selection historically. Yet, despite the striking repeatability in the mode of adaptation, the specific genes and functional pathways involved in cold compensation showed little overlap across genetic backgrounds. This implies that while the direction of selection on gene expression levels is predictable, the molecular routes to local adaptation are highly contingent on the evolutionary history of populations. One possible explanation for this low repeatability is that cold adaptation progresses through selection acting on exposed maladaptive cryptic genetic variation in gene expression that has accumulated randomly across the transcriptome on the different backgrounds (19,82–86). This randomness seems further mirrored in the low kME scores and gene co-regulation of cold compensation genes, suggesting that they tend to be peripheral in gene networks.

In contrast, the mode of heat adaptation was less consistent, comprising a mixture of genetic compensation and reinforcement. This finding echoes that of previous studies in mice (57) and fruit flies (51) reporting genetic reinforcement to be a relatively common mode of transcriptome adaptation to heat (but see studies on Anolis lizards (47) and flour beetles (49) for examples of more pronounced heat compensation). The relative prevalence of reinforcement suggests that plastic responses to heat can often be adaptive. In support of this hypothesis, heat adaptation in our data showed substantially greater convergence at the level of genes and biological functions compared to cold adaptation, although this was true for both reinforcement and compensation. Heat reinforcement genes also showed the highest kME values and co-regulation patterns in network analyses, suggesting that some of these may be pleiotropic hub genes that are repeatedly reused in evolutionary responses to mediate coordinated adaptive plasticity to changing environments (59,87,88).

An interesting result was the comparatively high repeatability of heat compensation genes and their annotated biological processes, compared to the low repeatability found for cold compensation genes. One possible explanation for this difference is if the heat compensation response in fact represents adaptive plasticity in form of an initial programmed and highly general stress response. Such a stress response, if costly, could become maladaptive in later stages of local adaptation, once the organism has achieved enough allelic turnover at other loci, and could with enough time result in a pattern of compensation. This adaptive scenario aligns well with the study by Rodríguez-Verdugo and colleagues (2016) who showed that patterns of heat compensation at several hundreds of initially over-expressed genes in *Escherichia coli* were due to only three adaptive mutations at a single gene encoding an RNA polymerase subunit that restored gene expression to ancestral (pre-stressed) levels through improved transcriptional efficiency (53) (see also: (49)). Testing if a similar process could explain the observed patterns in *C. maculatus* would ultimately require more fine-grained time- series data throughout our experiment. The greater gene co-regulation and higher repeatability of several biological processes associated with heat (compared to cold) compensation genes, of which many are “usual suspects” in the heat stress response (e.g. catalytic activity and responses to oxidative stress), aligns well with the hypothesis.

### The underlying genetic basis of reinforcement and compensation in thermal adaptation

If standing genetic variation is available, gene expression evolution via trans-regulatory loci is predicted to be rapid and more repeatable than polygenic adaptation through cis-regulatory changes at single genes (89,90). On the other hand, a lack of standing variation at major effect loci may cause high costs of adaptation if populations need to wait for the right allele to arrive by mutation (91,92), an effect that is exacerbated in the likely presence of deleterious pleiotropy (93–96). In the presence of genetic redundancy, polygenic responses via cis-regulatory loci may then offer a more viable route to adaptation and leave a more repeatable signal at the phenotype level, despite being idiosyncratic at the level of single nucleotide polymorphisms (97–102). While we detected directional selection within and upstream of coding regions of compensation and reinforcement genes, there was limited evidence that these changes were driving the observed expression changes. Although our analysis does not rule out that cis-regulation can play an important part in the expression evolution of a limited set of the studied loci, our findings support the view that transcriptome adaptation is primarily driven by trans-regulatory changes, as found for yeast, fruit flies and mice adapting to heat (57,88,103). However, if cis-regulatory evolution progressed through purifying selection on maladaptive alleles segregating at already low frequency (86–88,90), our analysis of directional selection based on allele frequency changes in pool-seq data from only two time-points some 60 generations apart would not have had the statistical power to separate such selected loci from those evolving under drift. Hence, we might have underestimated the amount of adaptive change at cis-regulatory loci.

Despite the low repeatability of expression evolution, candidate SNPs in and upstream of cold compensation genes showed the greatest repeatability across genetic backgrounds, and in contrast to the other three gene categories, this repeatability increased when the analysis was restricted to the few cold compensation genes that did show parallel expression evolution across backgrounds (Figure 7A). This indicates that, while allele frequency changes in differentially expressed genes were not associated with changes in their expression (Figure 7B, C), a correlation between sequence and expression evolution still exists. Such a correlation could be driven by some genes being more important than others (e.g. selection at cold temperature acts more strongly on both regulation and transcriptional efficacy of a few of key genes (31,104)), but further investigation would need to target why this pattern only was observed for cold compensation.

### Evolutionary repeatability at different levels of the genotype-phenotype map

The pattern we report here at the expression level - cold adaptation being predictable in direction but not in gene usage, and heat adaptation showing the reverse pattern - runs counter to what was previously observed at the level of sequence evolution in these experimental evolution lines (69), with cold adaptation showing greater gene usage across genetic backgrounds (see also Figure 7A). Nevertheless, at the level of fitness-related life history traits, heat adaptation again shows higher repeatability (69,70), which is more consistent with the current results we found in gene expression traits. The pattern for heat adaptation may not come as a big surprise; given a polygenic basis of adaptation, this result represents a common finding - there are likely many alternative genomic routes to achieve the same outcome at the level of locally adapted phenotypes. Our results on the repeatability of heat adaptation at the three hierarchical levels of the genotype-phenotype map (coding sequences; expression levels; life-history phenotypes) also align well with gene expression traits representing an intermediate between genotype and phenotype. What seems more surprising is the relatively low repeatability of cold adaptation at the gene expression level, coupled with its high repeatability at the sequence level. One possibility is that most of the observed repeatability at the sequence level encodes life history traits and gene expression in juveniles, while life history and gene expression data have only been collected from adult beetles so far (69). Juveniles are likely under strong selection at cold temperature due to intraspecific competition over limited food resources favouring fast growth and development while thermal constraints are strong on these traits (21,105,106). Future studies measuring gene expression and selection in juveniles under settings including competition might shed further light on this question.

### Conclusions

By combining replicated experimental evolution with both transcriptomic and pool-seq data, we explored whether thermal plasticity in gene expression is reinforced or compensated, and whether these evolutionary responses are repeatable across populations. We also asked if responses are underpinned by shared or distinct genetic architectures and biological functions. Our findings show that cold adaptation proceeded via many molecular routes that converged on a common compensatory outcome, suggesting that ancestral transcriptome plasticity to cold temperature is often maladaptive and corrected back to ancestral levels through genetic adaptation. Heat adaptation was more repeatable at the molecular level but less consistent in its direction relative to ancestral plasticity, encompassing almost equal amounts of compensation and reinforcement. We found reinforcement genes to occupy more central positions in gene networks and to be used repeatedly in the evolutionary responses, showing that the three genetic backgrounds all harbour genetic variation at loci regulating ancestral adaptive plasticity to heat. Yet, despite detecting signals of selection on DNA sequences of differentially expressed genes, these signals did not predict expression evolution, and repeatability of sequence evolution was poorly aligned with repeatability of expression changes. Thus, evolutionary repeatability in hot and cold climates may critically depend on the level of biological organization at which repeatability is studied. Thus, when predicting how populations will respond to ongoing climate change, evolutionary repeatability and genetic constraints need to be studied on, and assigned to, specific levels of the genotype-phenotype map.

## Acknowledgements

This work was supported by grant no. 2022-01117 from Formas, grant no. CTS22:2101 from Carl Tryggers Stiftelse, and a grant from Kungliga Fysiografiska Sällskapet i Lund to D.B. Sequencing was performed by the SNP&SEQ Technology Platform in Uppsala. The facility is part of the National Genomics Infrastructure (NGI) Sweden and Science for Life Laboratory. The SNP&SEQ Platform is also supported by the Swedish Research Council and the Knut and Alice Wallenberg Foundation. We would like to thank Milena Trabert for sharing her illustrations of *C. maculatus*, used in figure 1, and Johanna Liljestrand-Rönn för help in the lab.

