## Supplementary Information for "The mode and repeatability of thermal adaptation in gene expression in the seed beetle *Callosobruchus maculatus*"

**These authors contributed equally*

### Phenotypic Adaptation

To quantify adaptation in life-history phenotypes at the three assay temperatures, we re-analyzed data from von Schmalensee et al. (2026)[1]. These data were taken from all lines for juvenile development rate and adult reproduction using the siblings of the individuals measured for gene expression. Adult reproduction was assayed by placing a newly emerged (0-24h old) virgin male and female together in a 60mm petri-dish with *ad libitum* black-eyed beans. The couple was then allowed to mate and females to lay eggs freely until their death. In total, 1042 assays were set up over the two years and three assay temperatures. As a measure of juvenile development rate, the time it took from when the couples were given beans until the first adult offspring was found was noted in each of the 597 assays that were set up in 2023.

We analysed data in linear mixed effect models using the lme4 package[2] for R[3]. Adult reproduction was analysed with offspring number as a Poisson distributed response, and juvenile development rate as a normally distributed response following log-transformation. For both models, evolution regime, assay temperature, and genetic background were added as crossed fixed effects, and year as an additional blocking factor. Line replicates nested within backgrounds were added as crossed random effects. Ancestral populations were not included in statistical analysis of the phenotype data as they were only represented by single populations but are shown graphically in Fig. 1B.

Assay temperature sped up development rate ($X^2$ = 2606.6, df = 2, P < 0.001, n = 597) but decreased adult reproduction ($X^2$ = 72.6, df = 2, P < 0.001, n = 1042) (Fig. 1B). Hot lines developed slower ($X^2$ = 24.3, df = 1, P = 0.0026) but produced more offspring ($X^2$ = 100.0, df = 2, P < 0.001) than cold lines in a pattern congruent with thermal compensation (countergradient variation) (Fig. 1B, Supplementary Tables 1A and 1B). Only offspring production showed signs of temperature-specific adaptation (regime:temperature: $X^2$ = 21.8, df = 2, P < 0.001), with heat-adapted lines outperforming cold-adapted lines most readily at hot temperature, but surprisingly also more at cold temperature compared to benign 29°C (Fig. 1B). There were weak, but marginally (non)significant three-way interactions between regime, assay temperature and genetic background for both development rate ($X^2$ = 2.73, df = 4, P = 0.08) and offspring production ($X^2$ = 9.54, df = 4, P = 0.049), suggesting that thermal adaptation in life-history depended on the genetic background on which it occurred (Fig. 1B, Supplementary Tables 1a and 1B). These results replicate estimates taken at earlier stages of experimental evolution[4,5].

#### Supplementary Table 1a: Adult Reproduction

Mixed effect model on the effects of evolution regime (Hot vs. Cold lines) and assay temperature (23, 29 & 35C) on offspring production (log(number of offspring+1). Note that the ancestral populations were removed from the analysis since they are only represented by a single replicate and hence are not replicated in the strictest sense.

```
Generalized linear mixed model fit by maximum likelihood (Laplace Approximation) ['glmerMod']
 Family: poisson  ( log )
Formula: OF - 2 ~ regime * temperature * background + Year + (1 | replicate) +      (1 | temperature:replicate)
   Data: growthrates2023[growthrates2023$regime != "Ancestor", ]

      AIC       BIC    logLik -2*log(L)  df.resid
  12725.8   12829.7   -6341.9   12683.8      1021

Scaled residuals:
   Min     1Q Median     3Q    Max
-8.573 -1.002  0.258  1.318  5.981

Random effects:
 Groups                Name        Variance  Std.Dev.
 temperature:replicate (Intercept) 0.0042189 0.06495
 replicate             (Intercept) 0.0003056 0.01748
Number of obs: 1042, groups:  temperature:replicate, 36; replicate, 12
```

```
Analysis of Deviance Table (Type II Wald chisquare tests)

Response: OF - 2
                               Chisq Df Pr(>Chisq)
regime                       99.9768  1  < 2.2e-16 ***
temperature                  72.5766  2  < 2.2e-16 ***
background                    3.3952  2    0.18312
Year                          1.0931  1    0.29577
regime:temperature           21.7521  2  1.891e-05 ***
regime:background             1.1443  2    0.56432
temperature:background        5.2221  4    0.26526
regime:temperature:background 9.5395  4    0.04894 *
```

#### Supplementary Table 1b: Juvenile Development

Mixed effect model on the effects of evolution regime (Hot vs. Cold lines) and assay temperature (23, 29 & 35C) on development time (log(days of development from egg to adult)). Note that the ancestral populations were removed from the analysis since they are only represented by a single replicate and hence are not replicated in the strictest sense.

```
Linear mixed model fit by REML ['lmerMod']
Formula: log(Dev) ~ regime * temperature * background + (1 | replicate) +      (1 | temperature:replicate)
   Data: growthrates2023[growthrates2023$regime != "Ancestor", ]

REML criterion at convergence: -2083.1

Scaled residuals:
    Min      1Q  Median      3Q     Max
-3.9875 -0.5157 -0.0827  0.5091  6.6059

Random effects:
 Groups                Name        Variance  Std.Dev.
 temperature:replicate (Intercept) 5.886e-04 0.024262
 replicate             (Intercept) 4.366e-05 0.006607
 Residual                          1.358e-03 0.036857
Number of obs: 597, groups:  temperature:replicate, 36; replicate, 12
```

```
Analysis of Deviance Table (Type II Wald F tests with Kenward-Roger df)

Response: log(Dev)
                                      F Df  Df.res  Pr(>F)
regime                          24.3387  1  6.0012 0.00262 **
temperature                   2606.5577  2 12.0593 < 2e-16 ***
background                       2.0919  2  5.9986 0.20454
regime:temperature               0.9170  2 12.0830 0.42584
regime:background                0.2098  2  6.0050 0.81644
temperature:background           0.3123  4 12.0784 0.86430
regime:temperature:background    2.7262  4 12.0967 0.07932 .
```

### Repeatability of gene expression changes

| PAIR (index) | Label in spreadsheets | Plastic Baseline | Plastic Treatment | Evolved Baseline | Evolved Treatment |
|---|---|---|---|---|---|
| 1 | LogFC | A.Bra.1.29 | A.Bra.1.23 | A.Bra.2.23 | C.Bra.1.23 |
| 2 | LogFC.1 | A.Bra.2.29 | A.Bra.2.23 | A.Bra.1.23 | C.Bra.2.23 |
| 3 | LogFC.2 | A.Bra.1.29 | A.Bra.1.35 | A.Bra.2.35 | H.Bra.1.35 |
| 4 | LogFC.3 | A.Bra.2.29 | A.Bra.2.35 | A.Bra.1.35 | H.Bra.2.35 |
| 5 | LogFC.4 | A.Cal.1.29 | A.Cal.1.23 | A.Cal.2.23 | C.Cal.1.23 |
| 6 | LogFC.5 | A.Cal.2.29 | A.Cal.2.23 | A.Cal.1.23 | C.Cal.2.23 |
| 7 | LogFC.6 | A.Cal.1.29 | A.Cal.1.35 | A.Cal.2.35 | H.Cal.1.35 |
| 8 | LogFC.7 | A.Cal.2.29 | A.Cal.2.35 | A.Cal.1.35 | H.Cal.2.35 |
| 9 | LogFC.8 | A.Yem.1.29 | A.Yem.1.23 | A.Yem.1.23.2 | C.Yem.1.23 |
| 10 | LogFC.9 | A.Yem.2.29 | A.Yem.1.23.2 | A.Yem.1.23 | C.Yem.2.23 |
| 11 | LogFC.10 | A.Yem.1.29 | A.Yem.1.35 | A.Yem.1.35.2 | H.Yem.1.35 |
| 12 | LogFC.11 | A.Yem.2.29 | A.Yem.1.35.2 | A.Yem.1.35 | H.Yem.2.35 |
| 13 | LogFC.12 | A.Bra.1.29.n | A.Bra.1.23.n | A.Bra.2.23.n | C.Bra.1.23.n |
| 14 | LogFC.13 | A.Bra.2.29.n | A.Bra.2.23.n | A.Bra.1.23.n | C.Bra.2.23.n |
| 15 | LogFC.14 | A.Bra.1.29.n | A.Bra.1.35.n | A.Bra.2.35.n | H.Bra.1.35.n |
| 16 | LogFC.15 | A.Bra.2.29.n | A.Bra.2.35.n | A.Bra.1.35.n | H.Bra.2.35.n |
| 17 | LogFC.16 | A.Cal.1.29.n | A.Cal.1.23.n | A.Cal.2.23.n | C.Cal.1.23.n |
| 18 | LogFC.17 | A.Cal.2.29.n | A.Cal.2.23.n | A.Cal.1.23.n | C.Cal.2.23.n |
| 19 | LogFC.18 | A.Cal.1.29.n | A.Cal.1.35.n | A.Cal.2.35.n | H.Cal.1.35.n |
| 20 | LogFC.19 | A.Cal.2.29.n | A.Cal.2.35.n | A.Cal.1.35.n | H.Cal.2.35.n |
| 21 | LogFC.20 | A.Yem.1.29.n | A.Yem.1.23.n | A.Yem.1.23.2.n | C.Yem.1.23.n |
| 22 | LogFC.21 | A.Yem.1.29.2.n | A.Yem.1.23.2.n | A.Yem.1.23.n | C.Yem.2.23.n |
| 23 | LogFC.22 | A.Yem.1.29.n | A.Yem.1.35.n | A.Yem.2.35.n | H.Yem.1.35.n |
| 24 | LogFC.23 | A.Yem.1.29.2.n | A.Yem.2.35.n | A.Yem.1.35.n | H.Yem.2.35.n |

**Supplementary Table 2.** Schematic showing the rotation of ancestral samples to produce independent estimates of the correlation between plasticity and genetic adaptation in gene expression (presented in Fig. 4C of the main text). Sample labels follow the convention *regime.origin.replicate.assay[.n][.2]*, where regime is A (ancestral), C (cold-evolved), or H (hot-evolved); origin is Yemen, Brazil, or California; assay is the assay temperature in °C; ".n" denotes 2023 sampling (absent for 2022); and a trailing ".2" denotes a re-sequenced 2022 sample.

**Supplementary Table 3:** Overlap of ancestral thermal plasticity in gene expression between the three genetic backgrounds according to the four gene categories; Antagonistic (expression levels responding in opposite directions for hot and cold assay temperature), Synergistic (response in the same direction), Heat-limited (response only to heat), and Cold-limited (response only to cold).

| Intersections | Degree | Gene Category | Observed overlap | Expected overlap | Obs/Exp | P value |
|---|---|---|---|---|---|---|
| Yemen | 1 | Antagonistic Responses | 785 | NA | NA | NA |
| USA | 1 | Antagonistic Responses | 121 | NA | NA | NA |
| USA & Yemen | 2 | Antagonistic Responses | 59 | 14.1 | 4.19 | 1.8e-24 |
| Brazil | 1 | Antagonistic Responses | 746 | NA | NA | NA |
| Brazil & Yemen | 2 | Antagonistic Responses | 352 | 86.9 | 4.05 | 4.6e-156 |
| Brazil & USA | 2 | Antagonistic Responses | 53 | 13.4 | 4.0 | 2.0e-20 |
| Brazil & USA & Yemen | 3 | Antagonistic Responses | 43 | 1.6 | 27.6 | 3.2e-50 |
| Yemen | 1 | Synergistics Responses | 187 | NA | NA | NA |
| USA | 1 | Synergistics Responses | 136 | NA | NA | NA |
| USA & Yemen | 2 | Synergistics Responses | 19 | 3.8 | 5.03 | 4.7e-9 |
| Brazil | 1 | Synergistics Responses | 237 | NA | NA | NA |
| Brazil & Yemen | 2 | Synergistics Responses | 11 | 6.6 | 1.67 | 0.06 |
| Brazil & USA | 2 | Synergistics Responses | 10 | 4.8 | 2.09 | 0.02 |
| Brazil & USA & Yemen | 3 | Synergistics Responses | 1 | 0.1 | 7.53 | 0.12 |
| Yemen | 1 | Heat-limited | 1831 | NA | NA | NA |
| USA | 1 | Heat-limited | 1399 | NA | NA | NA |
| USA & Yemen | 2 | Heat-limited | 522 | 380.1 | 1.37 | 5.3e-21 |
| Brazil | 1 | Heat-limited | 1981 | NA | NA | NA |
| Brazil & Yemen | 2 | Heat-limited | 822 | 538.2 | 1.53 | 1.3e-62 |
| Brazil & USA | 2 | Heat-limited | 646 | 411.3 | 1.57 | 3.3e-51 |
| Brazil & USA & Yemen | 3 | Heat-limited | 294 | 111.7 | 2.63 | 1.1e-59 |
| Yemen | 1 | Cold-limited | 761 | NA | NA | NA |
| USA | 1 | Cold-limited | 479 | NA | NA | NA |
| USA & Yemen | 2 | Cold-limited | 90 | 54.1 | 1.66 | 3.7e-7 |
| Brazil | 1 | Cold-limited | 1008 | NA | NA | NA |
| Brazil & Yemen | 2 | Cold-limited | 157 | 113.8 | 1.38 | 4.6e-6 |
| Brazil & USA | 2 | Cold-limited | 83 | 71.6 | 1.15 | 0.08 |
| Brazil & USA & Yemen | 3 | Cold-limited | 26 | 8.1 | 3.21 | 2.2e-7 |

**Supplementary Table 4:** Overlap of genetic responses in gene expression between the three genetic backgrounds according to the four gene categories; Antagonistic (expression levels responding in opposite directions for heat and cold adaptation), Synergistic (response in the same direction), Heat-limited (response only to heat), and Cold-limited (response only to cold).

| Intersections | Degree | Gene Category | Observed overlap | Expected overlap | Obs/Exp | P value |
|---|---|---|---|---|---|---|
| Yemen | 1 | Antagonistic Responses | 30 | NA | NA | NA |
| USA | 1 | Antagonistic Responses | 41 | NA | NA | NA |
| USA & Yemen | 2 | Antagonistic Responses | 0 | 0.018 | 0 | 1 |
| Brazil | 1 | Antagonistic Responses | 81 | NA | NA | NA |
| Brazil & Yemen | 2 | Antagonistic Responses | 0 | 0.36 | 0 | 1 |
| Brazil & USA | 2 | Antagonistic Responses | 1 | 0.49 | 2.02 | 0.39 |
| Brazil & USA & Yemen | 3 | Antagonistic Responses | 0 | 0.002 | 0 | 1 |
| Yemen | 1 | Synergistics Responses | 235 | NA | NA | NA |
| USA | 1 | Synergistics Responses | 274 | NA | NA | NA |
| USA & Yemen | 2 | Synergistics Responses | 13 | 9.6 | 1.36 | 0.16 |
| Brazil | 1 | Synergistics Responses | 144 | NA | NA | NA |
| Brazil & Yemen | 2 | Synergistics Responses | 15 | 5.0 | 2.99 | 0.0001 |
| Brazil & USA | 2 | Synergistics Responses | 10 | 5.9 | 1.71 | 0.07 |
| Brazil & USA & Yemen | 3 | Synergistics Responses | 1 | 0.2 | 4.90 | 0.18 |
| Yemen | 1 | Heat-limited | 694 | NA | NA | NA |
| USA | 1 | Heat-limited | 927 | NA | NA | NA |
| USA & Yemen | 2 | Heat-limited | 120 | 95.5 | 1.26 | 0.003 |
| Brazil | 1 | Heat-limited | 862 | NA | NA | NA |
| Brazil & Yemen | 2 | Heat-limited | 131 | 88.8 | 1.48 | 9.1e-7 |
| Brazil & USA | 2 | Heat-limited | 168 | 118.6 | 1.42 | 3.3e-7 |
| Brazil & USA & Yemen | 3 | Heat-limited | 27 | 12.2 | 2.21 | 0.0001 |
| Yemen | 1 | Cold-limited | 830 | NA | NA | NA |
| USA | 1 | Cold-limited | 925 | NA | NA | NA |
| USA & Yemen | 2 | Cold-limited | 139 | 113.9 | 1.22 | 0.005 |
| Brazil | 1 | Cold-limited | 935 | NA | NA | NA |
| Brazil & Yemen | 2 | Cold-limited | 138 | 115.2 | 1.20 | 0.009 |
| Brazil & USA | 2 | Cold-limited | 148 | 128.3 | 1.15 | 0.027 |
| Brazil & USA & Yemen | 3 | Cold-limited | 19 | 15.8 | 1.20 | 0.24 |

**Supplementary Table 5:** Overlap between the three genetic backgrounds in the genes used according to the four modes of gene expression adaptation; Cold Compensation (expression levels responding in opposite direction for cold plasticity and cold adaptation), Heat Compensation (expression levels responding in opposite direction for heat plasticity and heat adaptation), Cold Reinforcement (expression levels responding in same direction for cold plasticity and cold adaptation), Heat Reinforcement (expression levels responding in same direction for heat plasticity and heat adaptation).

| Intersections | Degree | Gene Category | Observed overlap | Expected overlap | Obs/Exp | P value |
|---|---|---|---|---|---|---|
| Yemen | 1 | Cold Compensation | 448 | NA | NA | NA |
| USA | 1 | Cold Compensation | 263 | NA | NA | NA |
| USA & Yemen | 2 | Cold Compensation | 26 | 17.5 | 1.49 | 0.026 |
| Brazil | 1 | Cold Compensation | 530 | NA | NA | NA |
| Brazil & Yemen | 2 | Cold Compensation | 33 | 35.2 | 0.94 | 0.68 |
| Brazil & USA | 2 | Cold Compensation | 21 | 20.7 | 1.02 | 0.51 |
| Brazil & USA & Yemen | 3 | Cold Compensation | 2 | 1.4 | 1.45 | 0.40 |
| Yemen | 1 | Heat Compensation | 352 | NA | NA | NA |
| USA | 1 | Heat Compensation | 278 | NA | NA | NA |
| USA & Yemen | 2 | Heat Compensation | 45 | 14.5 | 3.10 | 4.48e-12 |
| Brazil | 1 | Heat Compensation | 217 | NA | NA | NA |
| Brazil & Yemen | 2 | Heat Compensation | 35 | 11.3 | 3.09 | 1.4e-09 |
| Brazil & USA | 2 | Heat Compensation | 40 | 9.0 | 4.47 | 3.4e-16 |
| Brazil & USA & Yemen | 3 | Heat Compensation | 11 | 0.5 | 23.5 | 2.2e-12 |
| Yemen | 1 | Cold Reinforcement | 37 | NA | NA | NA |
| USA | 1 | Cold Reinforcement | 15 | NA | NA | NA |
| USA & Yemen | 2 | Cold Reinforcement | 1 | 0.1 | 12.1 | 0.08 |
| Brazil | 1 | Cold Reinforcement | 47 | NA | NA | NA |
| Brazil & Yemen | 2 | Cold Reinforcement | 7 | 0.3 | 27.1 | 4.4e-09 |
| Brazil & USA | 2 | Cold Reinforcement | 2 | 0.1 | 19.1 | 0.005 |
| Brazil & USA & Yemen | 3 | Cold Reinforcement | 1 | 0.0 | 1741 | 0.0006 |
| Yemen | 1 | Heat Reinforcement | 118 | NA | NA | NA |
| USA | 1 | Heat Reinforcement | 101 | NA | NA | NA |
| USA & Yemen | 2 | Heat Reinforcement | 8 | 1.8 | 4.52 | 0.0004 |
| Brazil | 1 | Heat Reinforcement | 278 | NA | NA | NA |
| Brazil & Yemen | 2 | Heat Reinforcement | 21 | 4.9 | 4.31 | 1.0e-08 |
| Brazil & USA | 2 | Heat Reinforcement | 21 | 4.2 | 5.04 | 5.2e-10 |
| Brazil & USA & Yemen | 3 | Heat Reinforcement | 3 | 0.1 | 41.1 | 5.8e-05 |

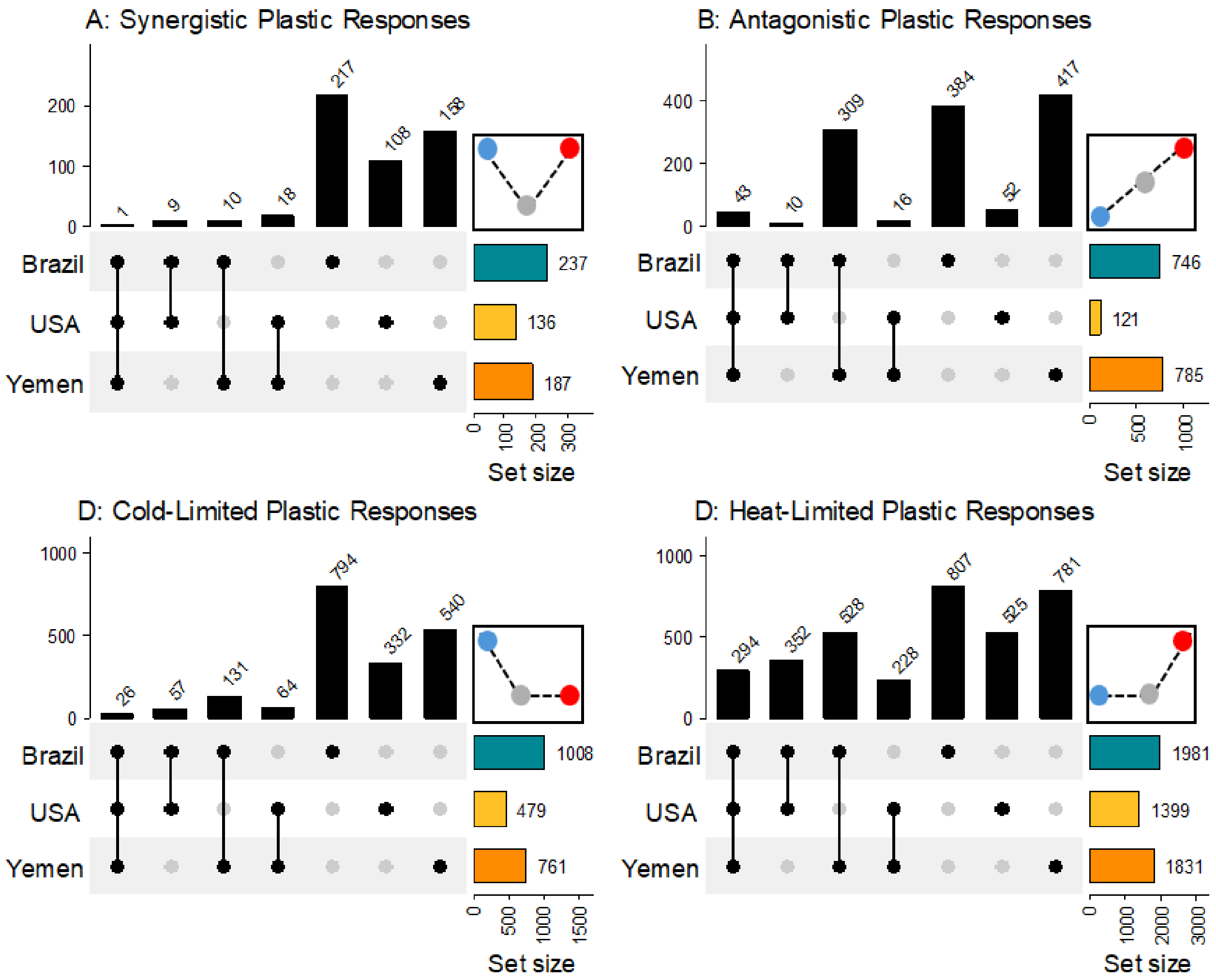


**Supplementary Figure 1: Ancestral thermal plasticity in gene expression**. The overlap between genetic backgrounds of differentially expressed genes in ancestors responding to temperature for each of the four gene classes (A: Synergistic, B: Antagonistic, C: Cold-limited, D: Heat-limited). Numbers above each bar represent the unique set of genes belonging to the category. Antagonistic, Cold-limited, and Heat-limited (but not Synergistic) genes all show much greater overlaps than expected by chance. A total of 6739 genes were analysed.

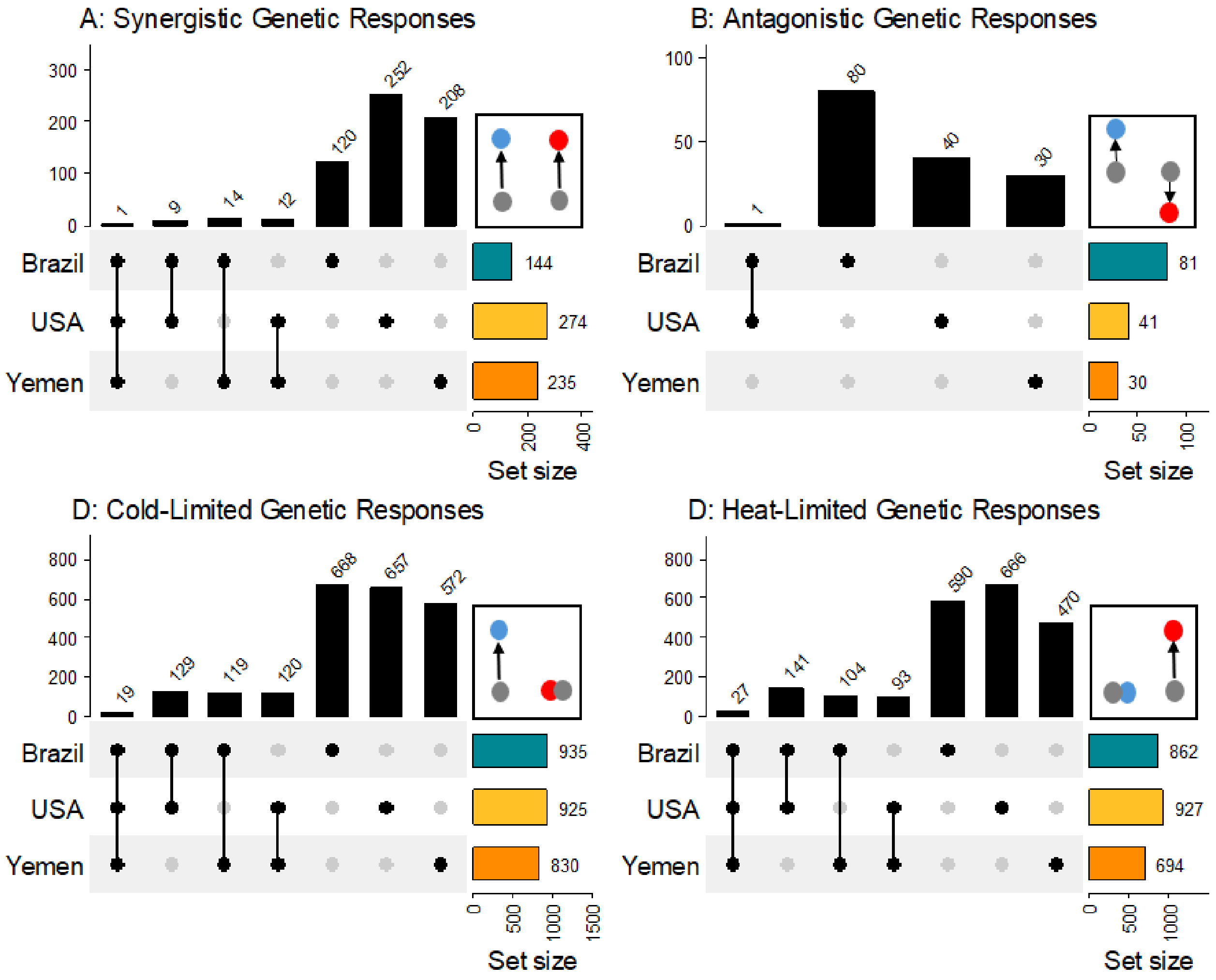


**Supplementary Figure 2**: **Thermal adaptation in gene expression.** The overlap between genetic backgrounds in differentially expressed genes due to genetic changes during experimental evolution for each of the four classes (A: Synergistic, B: Antagonistic, C: Cold-limited, D: Heat-limited). Numbers above each bar represent the unique set of genes belonging to the category. Cold-limited and Heat-limited genes show much greater overlaps than expected by chance whereas Antagonistic genes do not and Synergistic genes only marginally so. A total of 6739 genes were analysed.

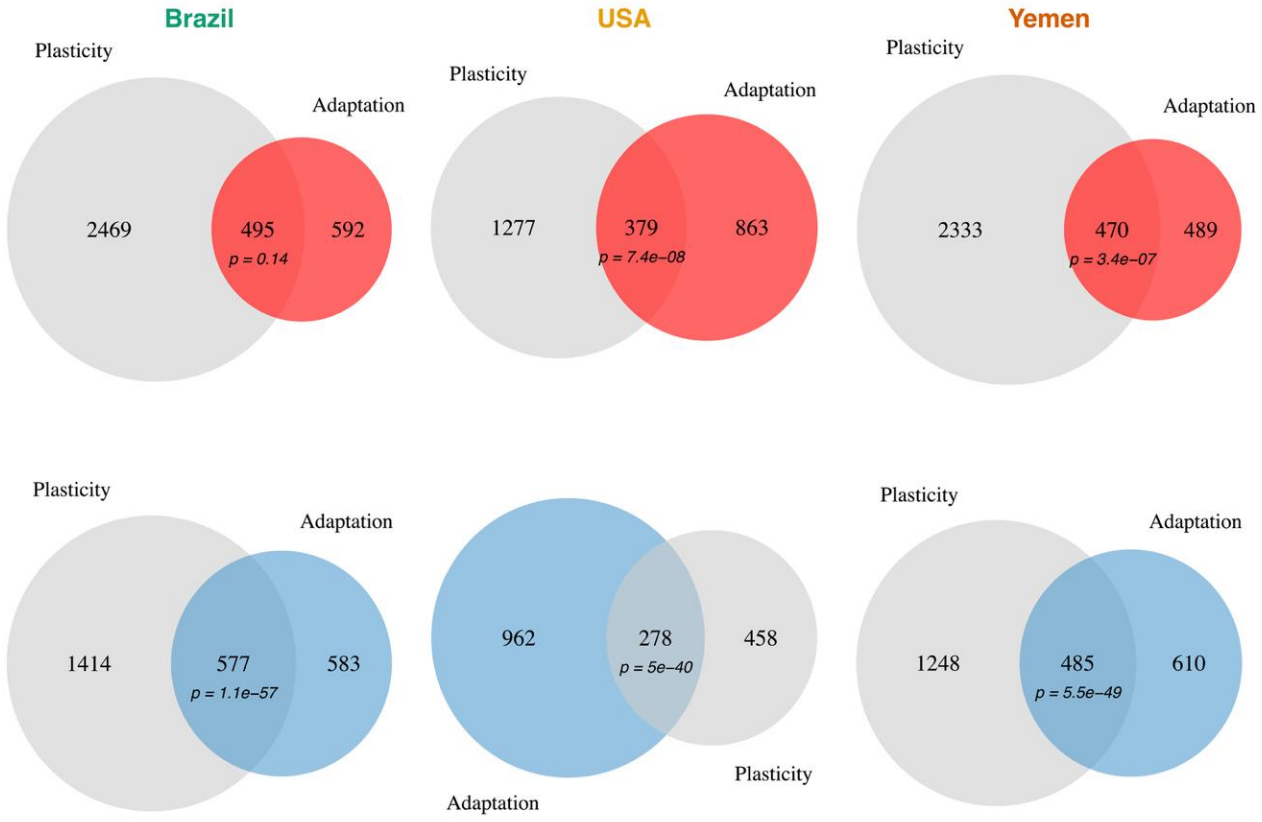


**Supplementary Figure 3:** The overlap between plastic (grey) and genetic (blue and red) responses was greater than expected by chance for both heat and cold, except for the Brazil background adapting to heat. However, the overlap was generally less pronounced for heat adaptation, with observed/expected ratios of; Brazil: 1.04, USA: 1.24, Yemen: 1.18, compared to cold adaptation; Brazil: 1.68, USA: 2.05, Yemen: 1.72.

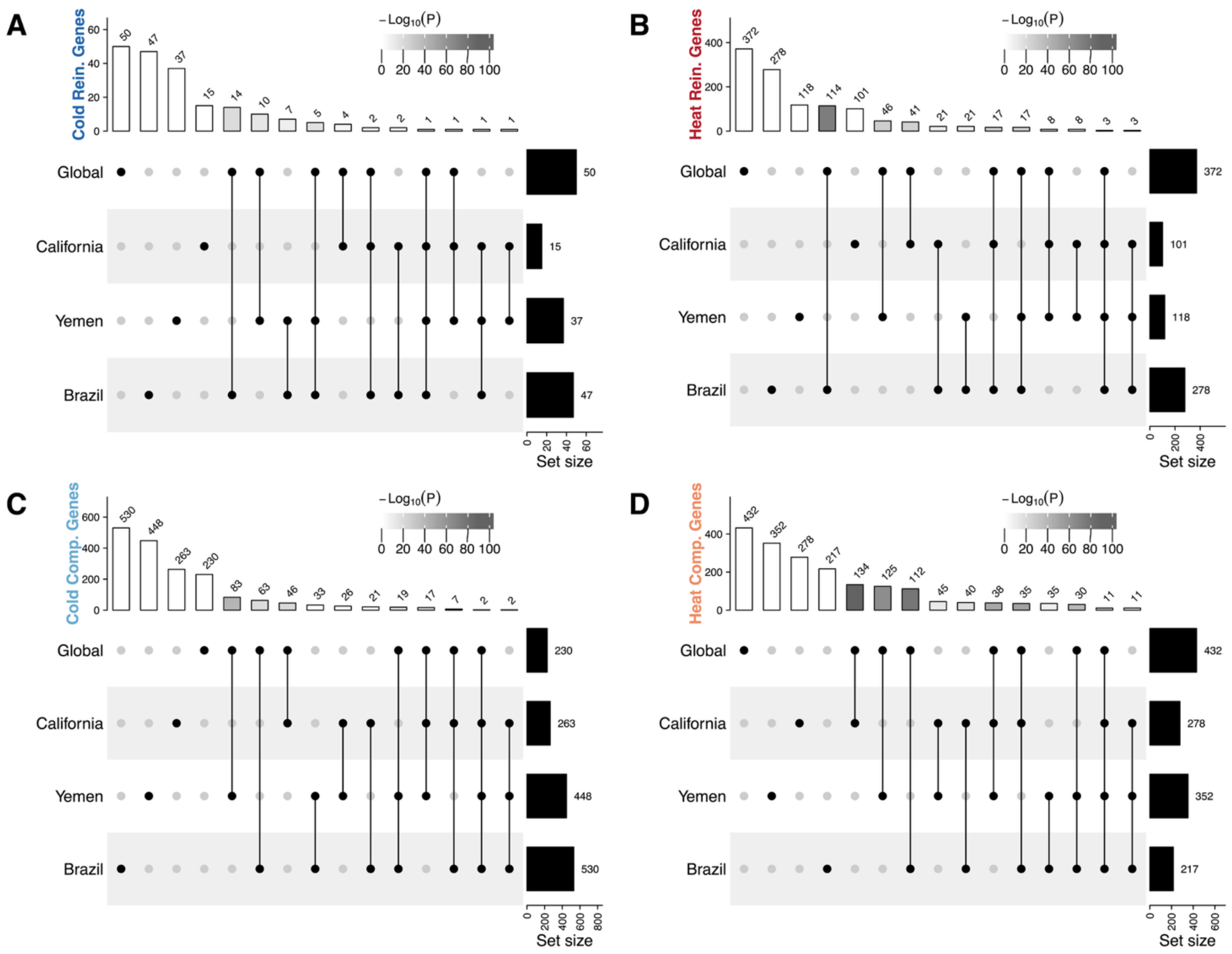


**Supplementary Figure 4**. **Overlap of compensation and reinforcement genes.** Do the backgrounds use the same genes when adapting through reinforcement (A, B) and compensation (C, D)? At hot temperature (B, D), both modes of adaptation show greater overlap than expected by chance. At cold temperature, however, only reinforcement shows greater overlap, whereas backgrounds do not share more genes involved in cold compensation than expected by chance (panel C). Moreover, while the global analysis picked up more significant genes than in the background-specific analyses for heat and cold reinforcement and heat compensation (as expected due to three times larger samples size), the global analysis for cold compensation picked up less significant genes than in any of the three background-specific analysis (panel C). Hence, while the pattern of cold compensation was the most pronounced and repeatable mode of adaptation, the involved genes are typically not shared across populations.

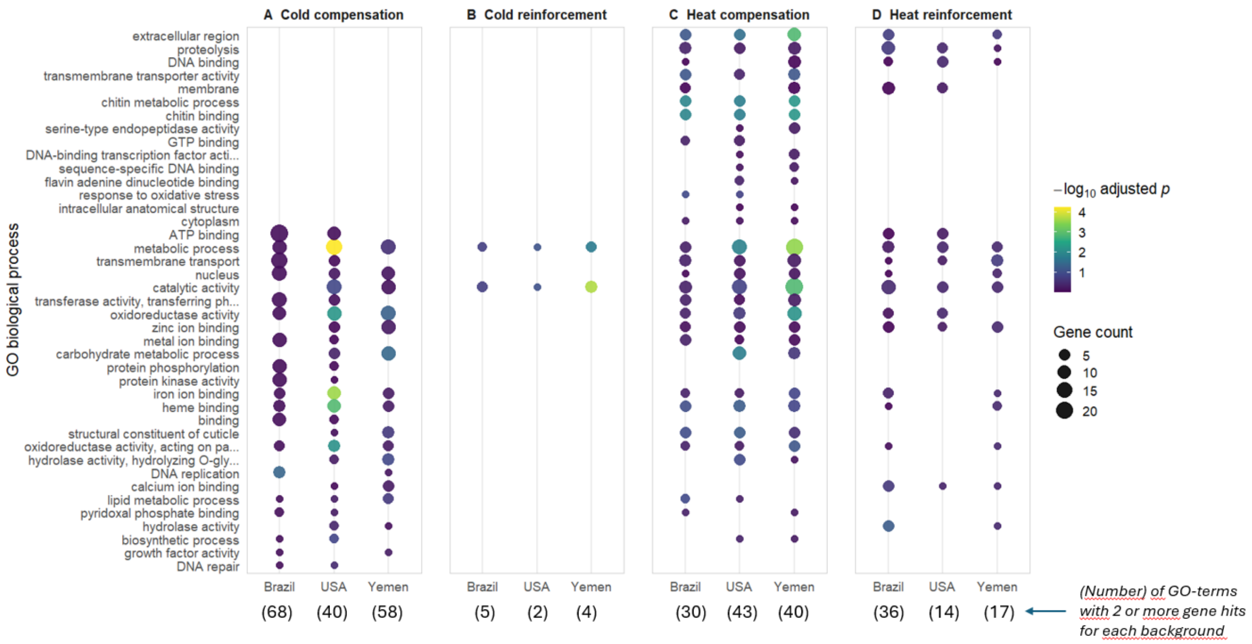


| | | | | |
|---|---|---|---|---|
| *Cut-off for included top-ranked terms (N)* N = 40 | P = 0.007 | P = 0.11 | P < 0.001 | P < 0.001 |
| N = 20 | P = 0.17 | P = 0.10 | P < 0.001 | P = 0.002 |
| N = 10 | P = 1 | P = 0.026 | P < 0.001 | P = 1 |

**Supplementary Figure 5**. **Overlap of GO-terms for different number of top-ranked terms.** Do the backgrounds adapt through the same gene functions (ontologies) for reinforcement (A, B) and compensation (C, D)? At hot temperature (B, D), both modes of adaptation show greater GO-term overlap than expected by chance except when gene lists are reduced to the top-10 ranked genes. At cold temperature, however, only reinforcement shows strong overlap, although the analysis is limited by the low number of cold reinforcement genes. Go overlap turns significant once GO-lists are increased to the top-40 ranked terms for cold compensation, suggesting that the same gene functions are to some extent targeted, but their relative importance is randomly distributed across the three backgrounds. Hence, while the pattern of cold compensation was the most pronounced and repeatable mode of adaptation, the involved gene functions do not show strong overlap across populations. P-values refer to the number of simulations in which permuted data show equal or greater fraction of overlapping terms compared to the observed data.

### Gene network analysis

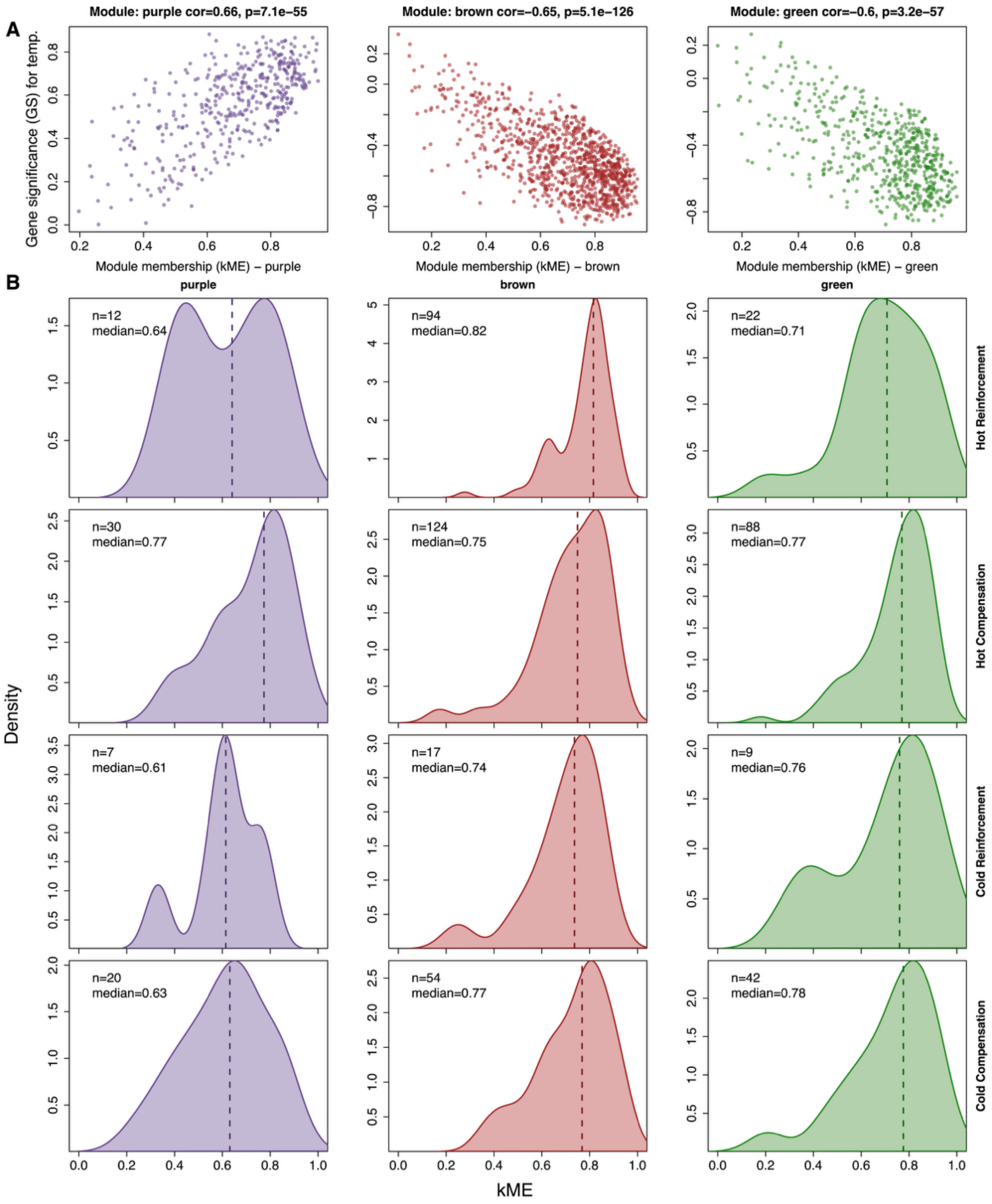


**Supplementary Figure 6. Network position of temperature-associated co-expression modules**. (A) Gene significance for temperature (GS, the correlation of each gene's expression with assay temperature) against module membership (kME, the correlation of each gene with its module eigengene) for the genes in each module. Each point is a gene; the panel header gives the module's GS–kME correlation and P-value. (B) Distribution of module membership (kME) for genes assigned to each module that also belong to each global compensation/reinforcement category. The dashed line marks the median kME; n is the number of genes in each module × category overlap. Higher kME indicates more hub-like (central) network position.

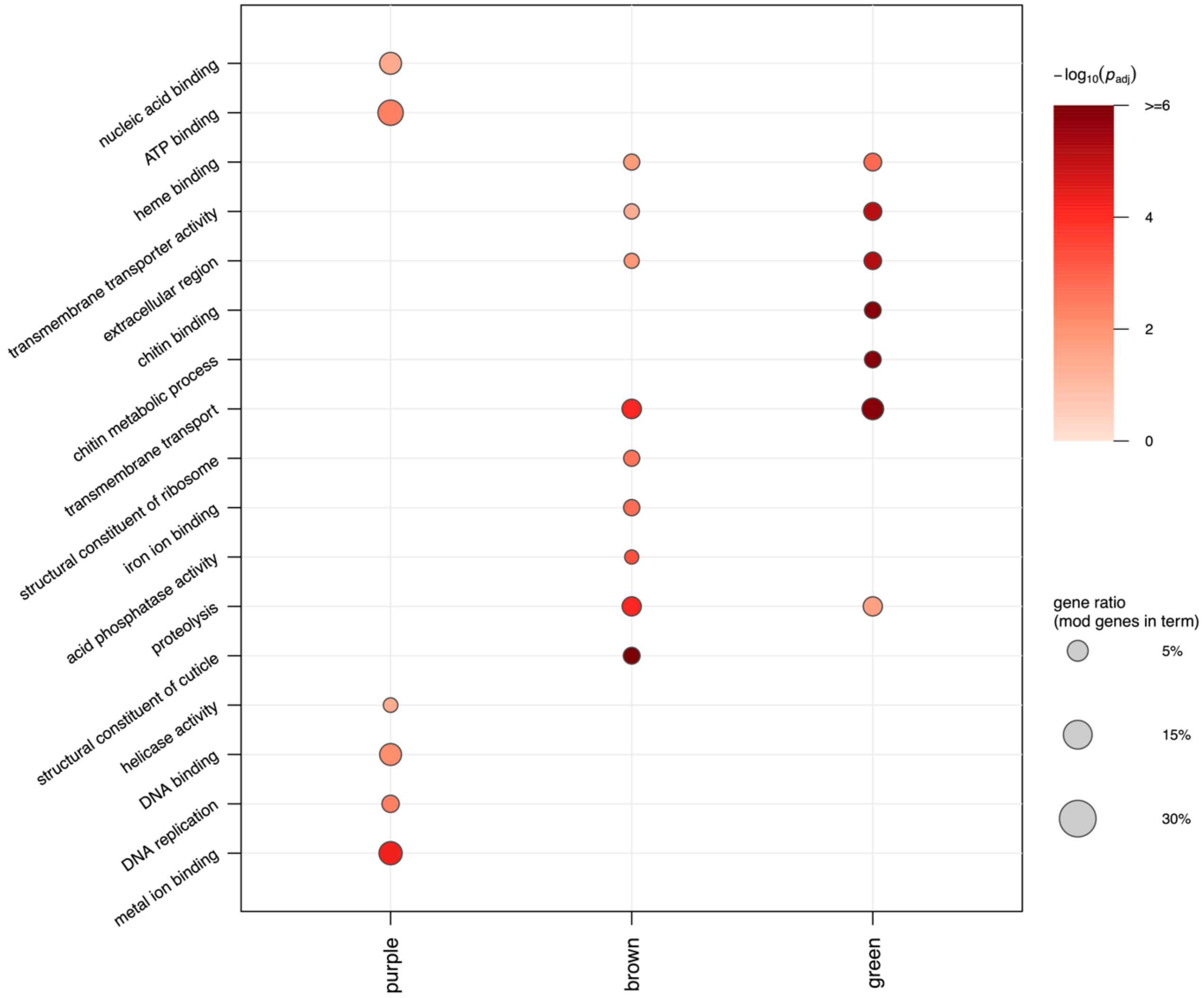


**Supplementary Figure 7: GO term over-representation for the three focal co-expression modules.** Per-module enrichment was tested with a hypergeometric test for each GO term using the genes in the WGCNA network as the background universe and Benjamini–Hochberg correction for multiple testing. For each module (purple, brown, green), the top six BH-significant terms ($p\_adj < 0.05$) are shown; a term appears in every module in which it was also significant. Dot color indicates significance and dot size the gene ratio.

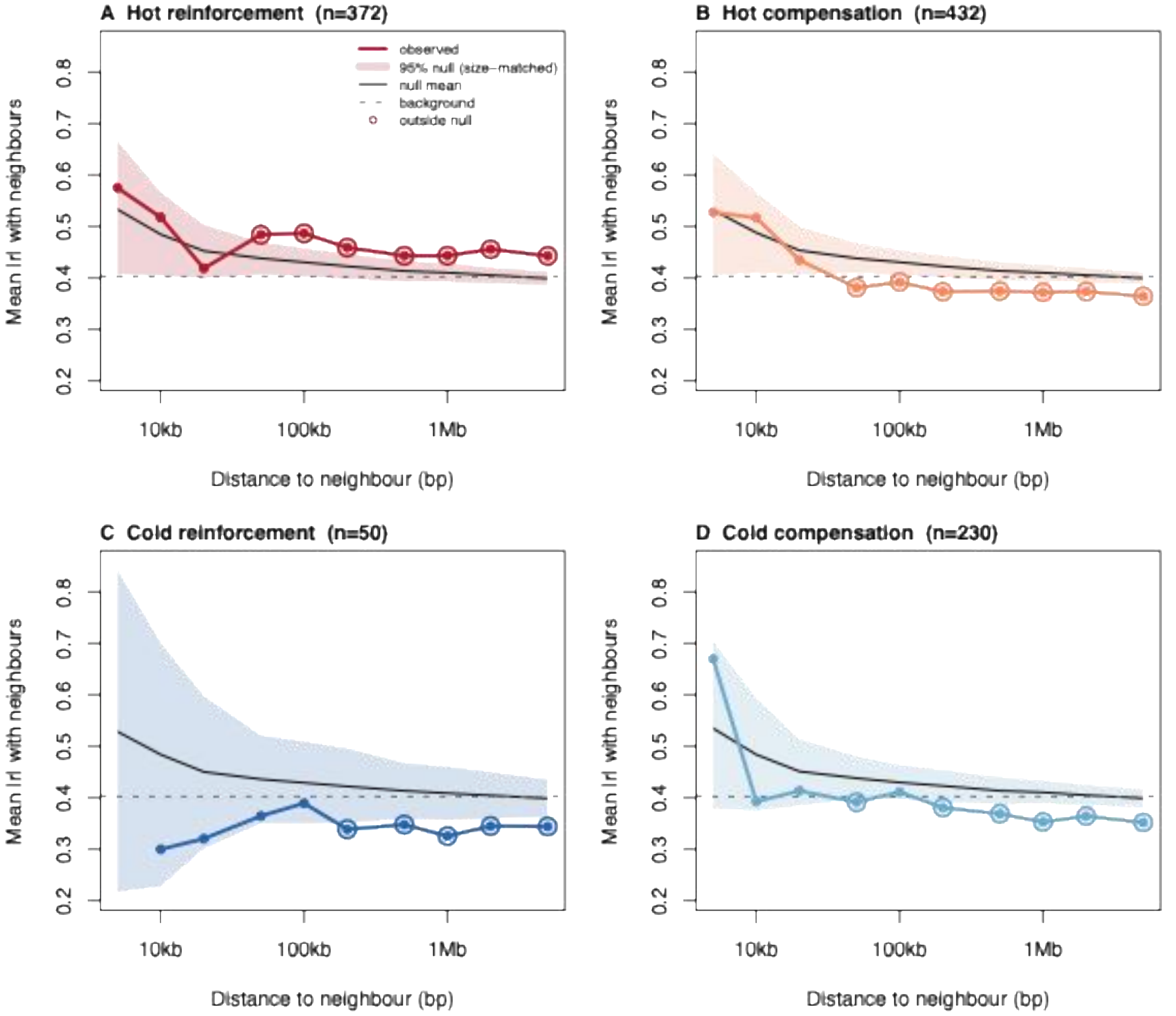


**Supplementary Figure 8:** Neighbourhood co-expression of each category against a size-matched null. Each panel shows the observed out-of-set neighbourhood decay for one gene category (A heat reinforcement, n = 372; B heat compensation, n = 432; C cold reinforcement, n = 50; D cold compensation, n = 230). Shaded band, bootstrapped 95% confidence interval from 1,000 size-matched random gene sets; solid black line, null mean; dashed grey line, cross-scaffold background; open circles, observed bins falling outside the 95% null. Heat reinforcement genes lie above the null at intermediate-to-large distances, whereas genes belonging to the other three categories lie at or below it.

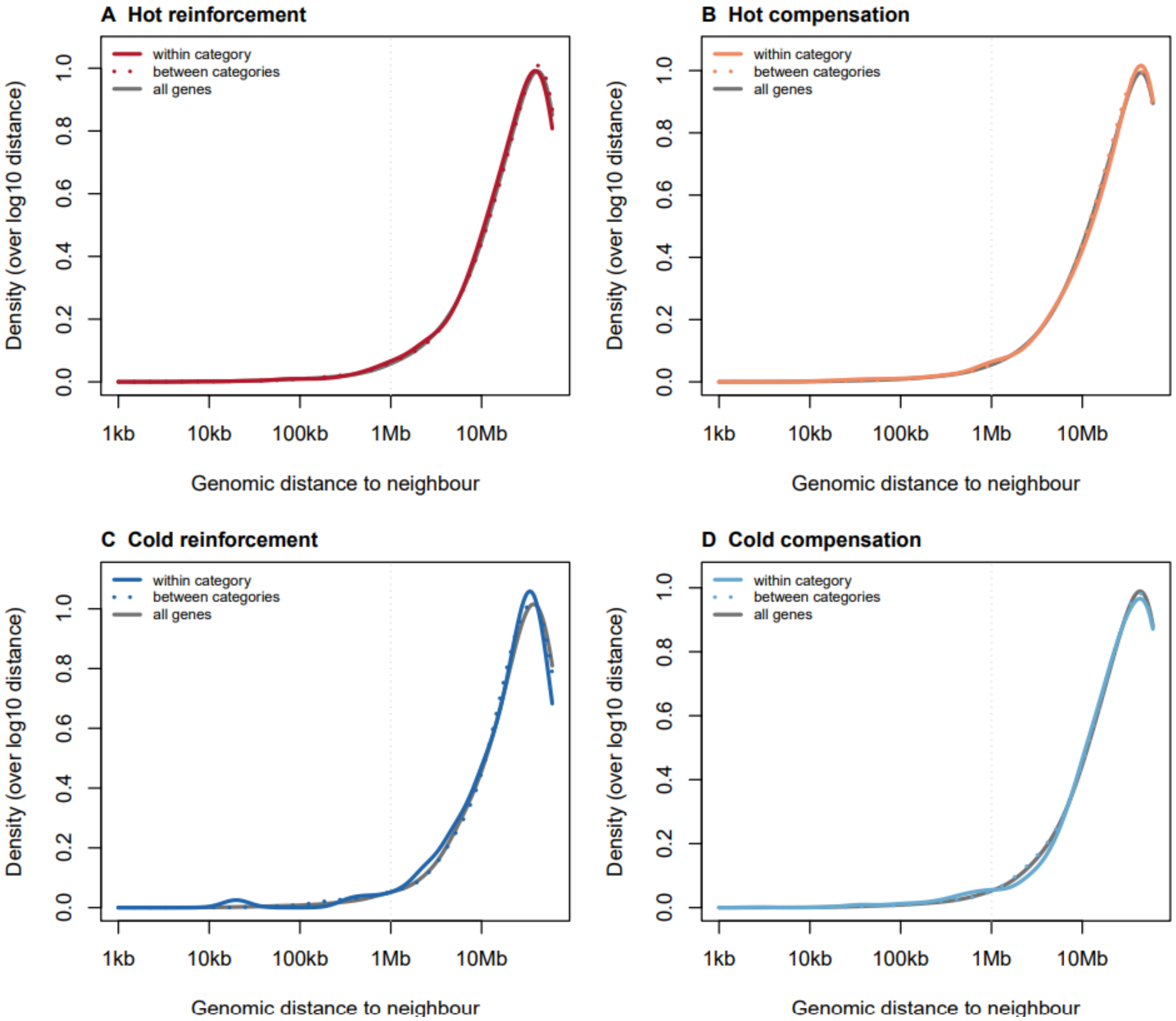


**Supplementary Figure 9: Pairwise genomic distance between focal genes and their neighbors.** Pairwise distances are shown between focal genes (belonging to one of the four adaptation categories) and other genes of either the same or different categories, as well as to all genes analyzed, on the same scaffold. No clear differences in physical distance exist to explain tighter co-regulation for heat reinforcement genes, suggesting trans-regulation as a causal mechanism.

### Estimates of purifying selection on DNA sequences of compensation and reinforcement genes.

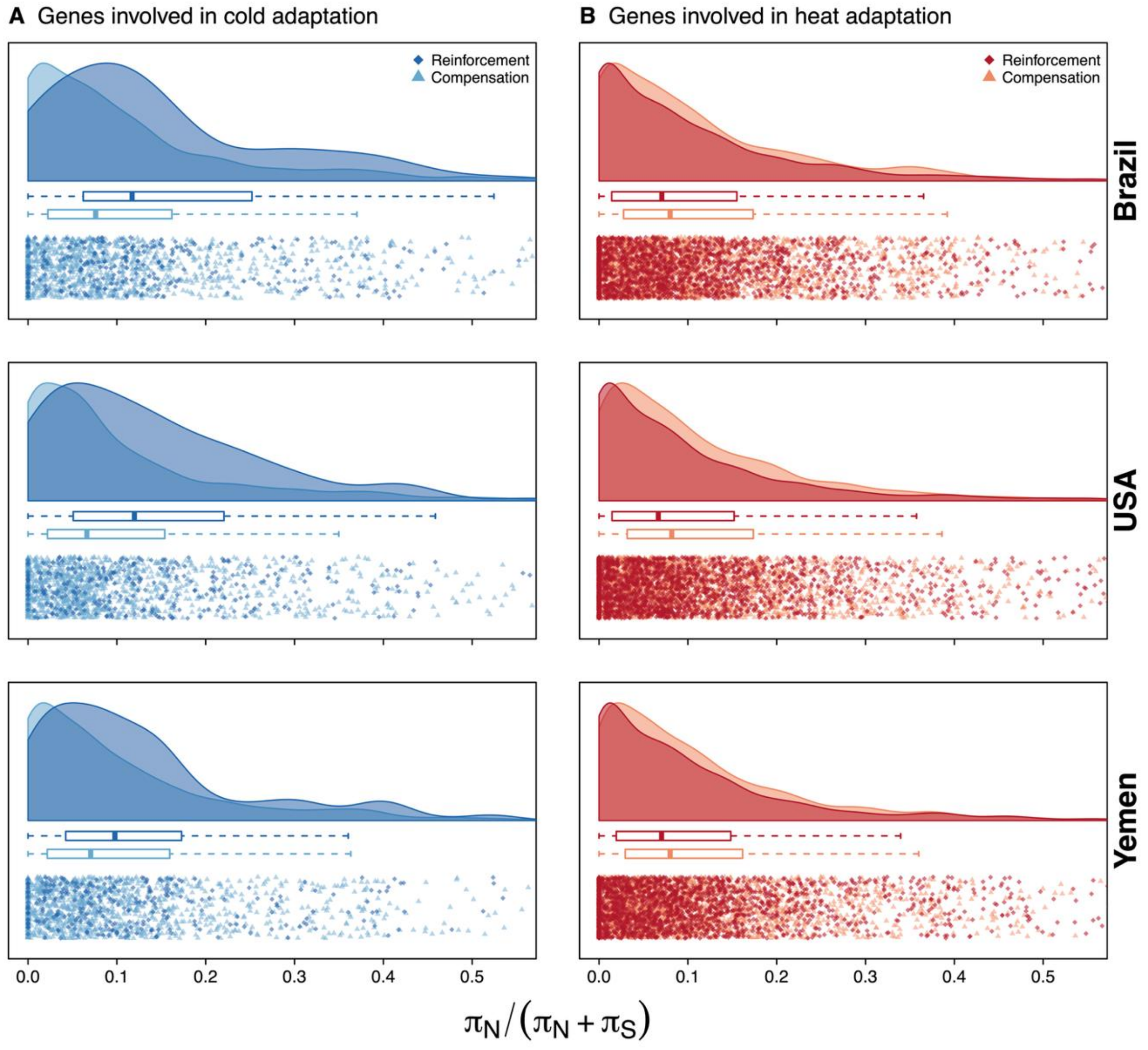


**Supplementary Figure 10: Purifying selection on compensation and reinforcement genes.** Genes involved in genetic reinforcement and compensation differ in their $\pi_N/(\pi_N+\pi_S)$ ratios, but the direction of the difference depends on the type of adaptation. For genes involved in heat adaptation (B), reinforcement genes have lower $\pi_N/(\pi_N+\pi_S)$ ratios than compensation genes, indicating historically stronger purifying selection on reinforcement genes. For genes involved in cold adaptation (A) the pattern is reversed, and larger in magnitude: reinforcement genes have higher $\pi_N/(\pi_N+\pi_S)$ ratios than compensation genes, indicating historically weaker purifying selection on reinforcement genes. Rows correspond to the three genetic backgrounds (Brazil, USA, Yemen). Density plots are based on 50 reinforcement and 230 compensation genes for cold adaptation, and 372 reinforcement and 432 compensation genes for heat adaptation. Each plot is based on six pool-seq samples per background.

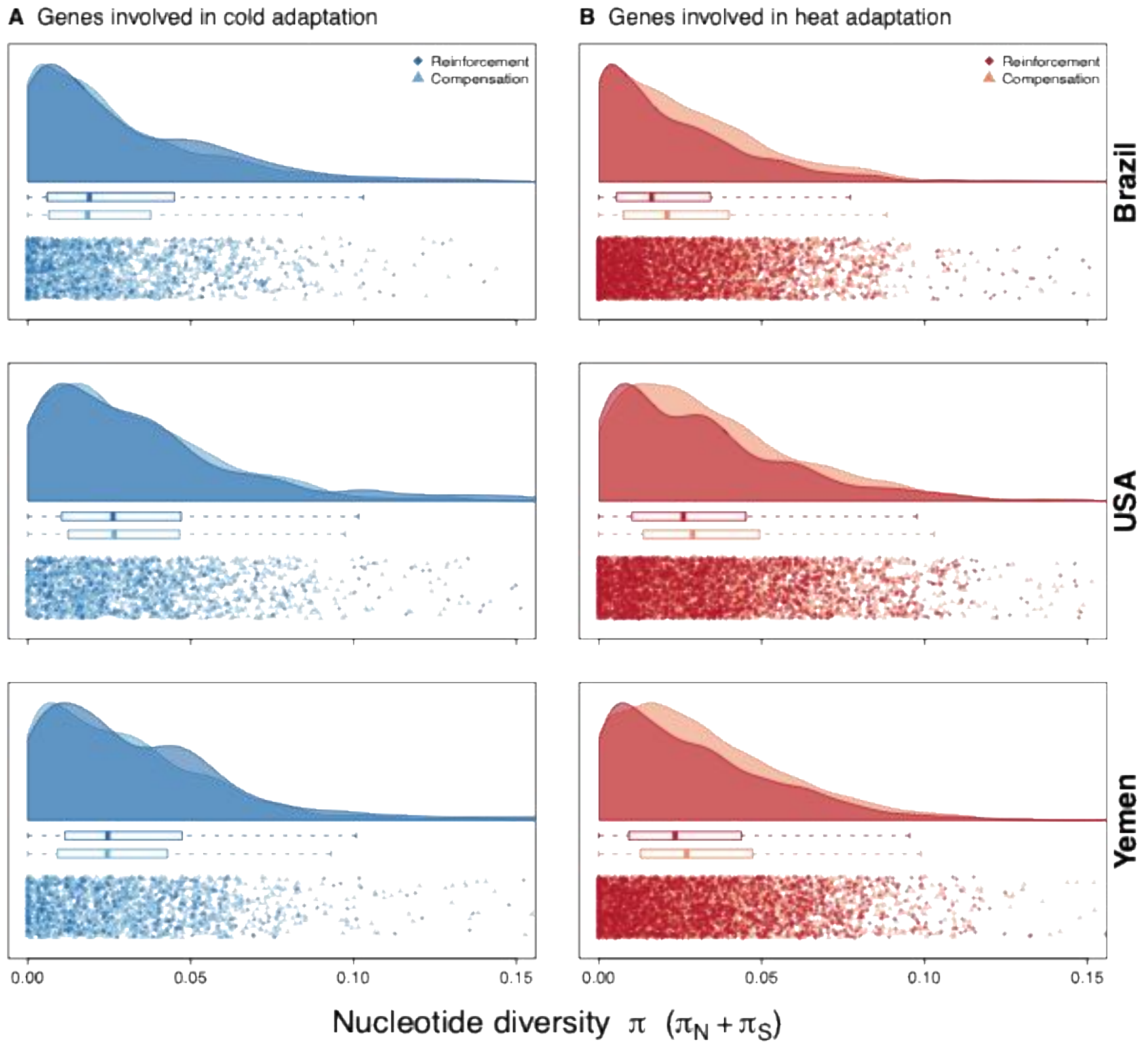


**Supplementary Figure 11: The amount of genetic diversity at cold and hot compensation and reinforcement genes.** For cold adaptation, genes involved in genetic compensation (light color) and genetic reinforcement (dark color) have similar levels of overall genetic diversity (pi). For heat adaptation, genes involved in genetic reinforcement have lower diversity on average. Density plots are based on 50 reinforcement and 230 compensation genes for cold adaptation, and 372 reinforcement and 432 compensation genes for heat adaptation. Each plot is based on pooling of the six-replicate pool-seq samples per background.

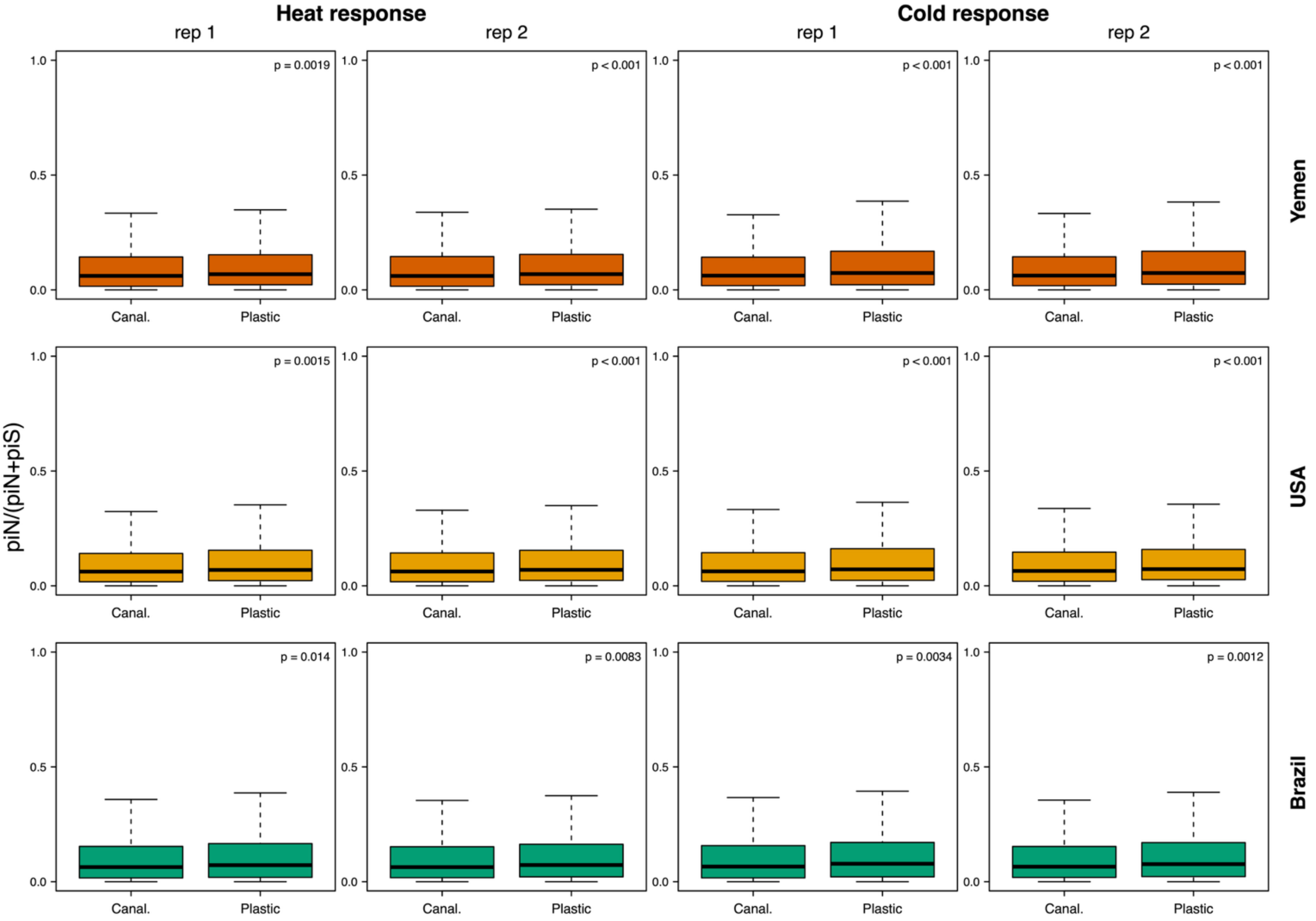


**Supplementary Figure 12:** For each ancestral pool-seq sample, the strength of purifying selection, estimated by the proportion of non-synonymous to total nucleotide polymorphisms $\pi_N/(\pi_N+\pi_S)$, is compared between genes that respond plastically to temperature and canalized (non-plastic) genes, for heat (left two columns) and cold (right two columns). Genes with a plastic expression response (i.e. conditional expression) have higher $\pi_N/(\pi_N+\pi_S)$ than canalized genes. A gene was classified as plastic if it changed expression significantly across the ancestral 29 to 35 °C (heat) or 29 to 23 °C (cold) contrast ($P \leq 0.05$); all other tested genes were considered canalized. Within each panel, $\pi_N/(\pi_N+\pi_S)$ is compared between the plastic and canalized gene sets using a two-sided Wilcoxon rank-sum; the test is unpaired and the unit of replication is the gene. In every ancestral sample and for both temperatures, plastic genes showed significantly higher $\pi_N/(\pi_N+\pi_S)$ than canalized genes, indicative of weaker purifying selection.

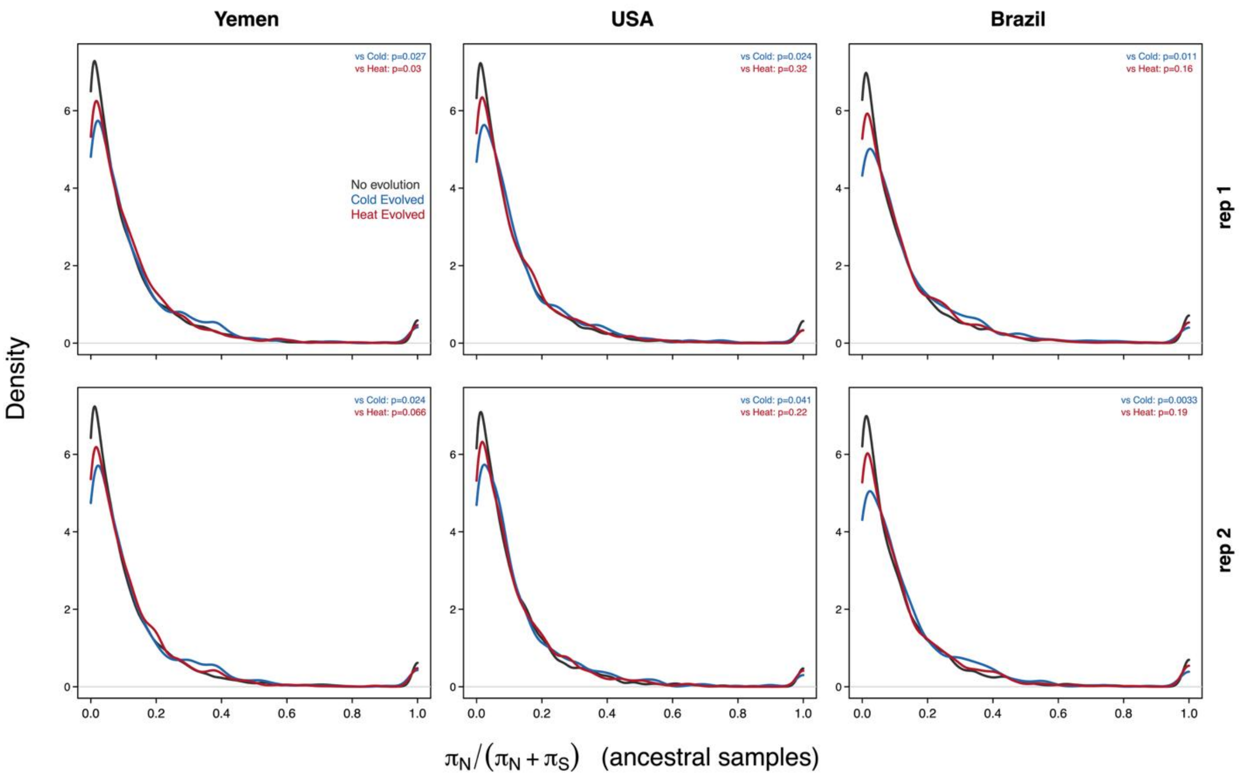


**Supplementary Figure 13:** For each ancestral pool-seq sample, the strength of purifying selection is compared between genes whose expression evolved during experimental evolution and genes whose expression did not, shown separately for cold-adaptation genes and heat-adaptation genes. A gene was classified as evolved if it showed a significant log-fold change in expression between the ancestor and the evolved line at each line's adaptation temperature; all other tested genes were classified as non-evolving. Within each panel, $\pi_N/(\pi_N+\pi_S)$ is compared between the evolved and non-evolving gene sets using a two-sided Wilcoxon rank-sum test; the test is unpaired and the unit of replication is the gene. Genes that did not evolve during experimental evolution were more likely to show low $\pi_N/(\pi_N+\pi_S)$ in ancestral samples, suggestive of being under stronger purifying selection relative to evolved genes, more so for genes involved in cold adaptation than those involved in heat adaptation. P-values are presented for Wilcoxon rank-sum tests of non-evolving genes vs cold- or heat-evolved genes.

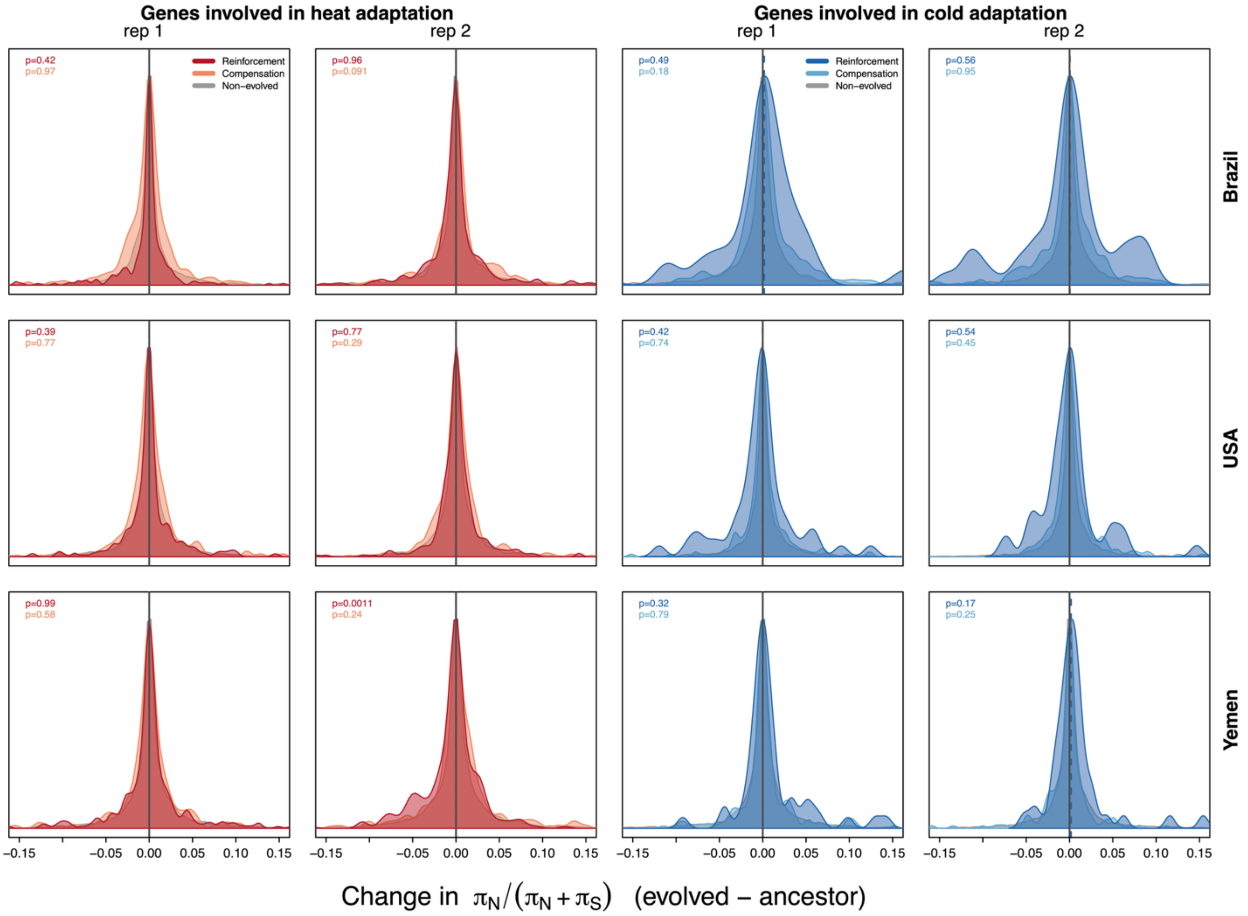


**Supplementary Figure 14:** For each origin's replicate lines, the change in the strength of purifying selection over experimental evolution (evolved – ancestor) is shown for genes involved in heat adaptation and cold adaptation. In each panel, the Δ distribution of reinforcement genes and of compensation genes (coloured curves) is compared against that of non-evolved genes (grey; genes with no significant evolved changes in expression) in the matched regime, which serve as the empirical null. The vertical line marks Δ = 0 and dashed lines mark category medians. Genes involved in heat and cold adaptation did not differ in how $\pi_N/(\pi_N+\pi_S)$ changed over the course of evolution relative to non-evolved genes, with the exception of Yemen replicate 2 for heat reinforcement, in which $\pi_N/(\pi_N+\pi_S)$ was slightly greater in the evolved than the ancestral line. Each panel reports two p-values, colour-coded by category, from two-sided Wilcoxon rank-sum (Mann–Whitney U) tests of reinforcement or compensation genes vs non-evolved genes; the tests are unpaired and the unit of replication is the gene.

### Estimates of directional selection on DNA sequences of compensation and reinforcement genes.

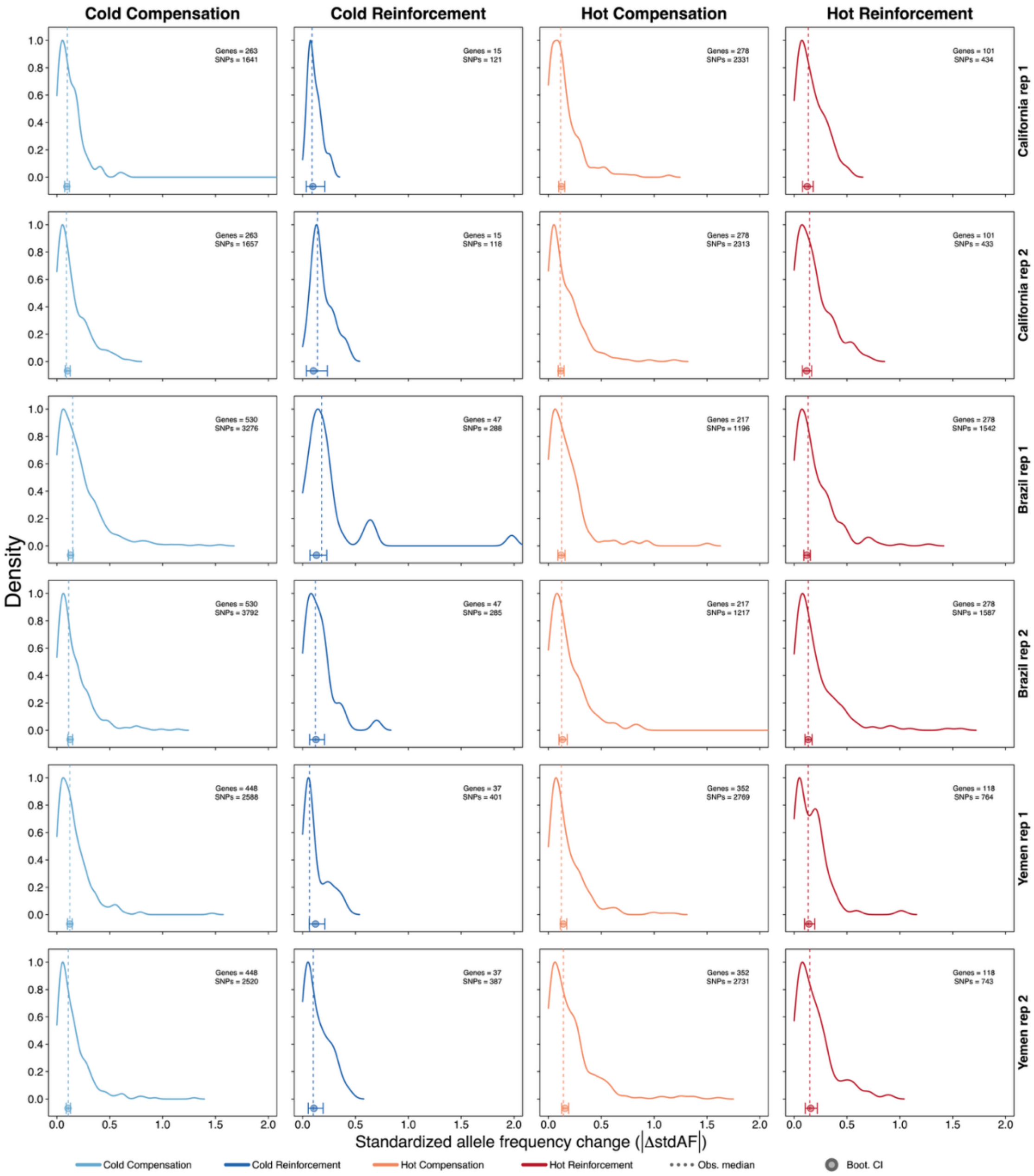


**Supplementary Figure 15: Standardized allele-frequency change in coding and 1kb-upstream regions of differentially expressed genes.** For each gene, the magnitude of standardized allele-frequency change was averaged across all non-synonymous SNPs in its gene body plus 1 kb upstream. Each panel shows the density of these per-gene mean values for each gene set category in each replicate population. Dotted vertical lines mark the observed median. Points and horizontal bars below each axis give the median and 95 % bootstrap interval expected under a background null (genes outside the focal category, resampled to the focal set size). For each bootstrap, *n* genes (matching the focal category size) were drawn with replacement from all genes outside that category in the same population and their median |ΔstdAF| recorded, repeated 2,000 times. Genes involved in genetic reinforcement or compensation do not show signs of being under stronger directional selection on average, although specific genes show signals of directional selection.

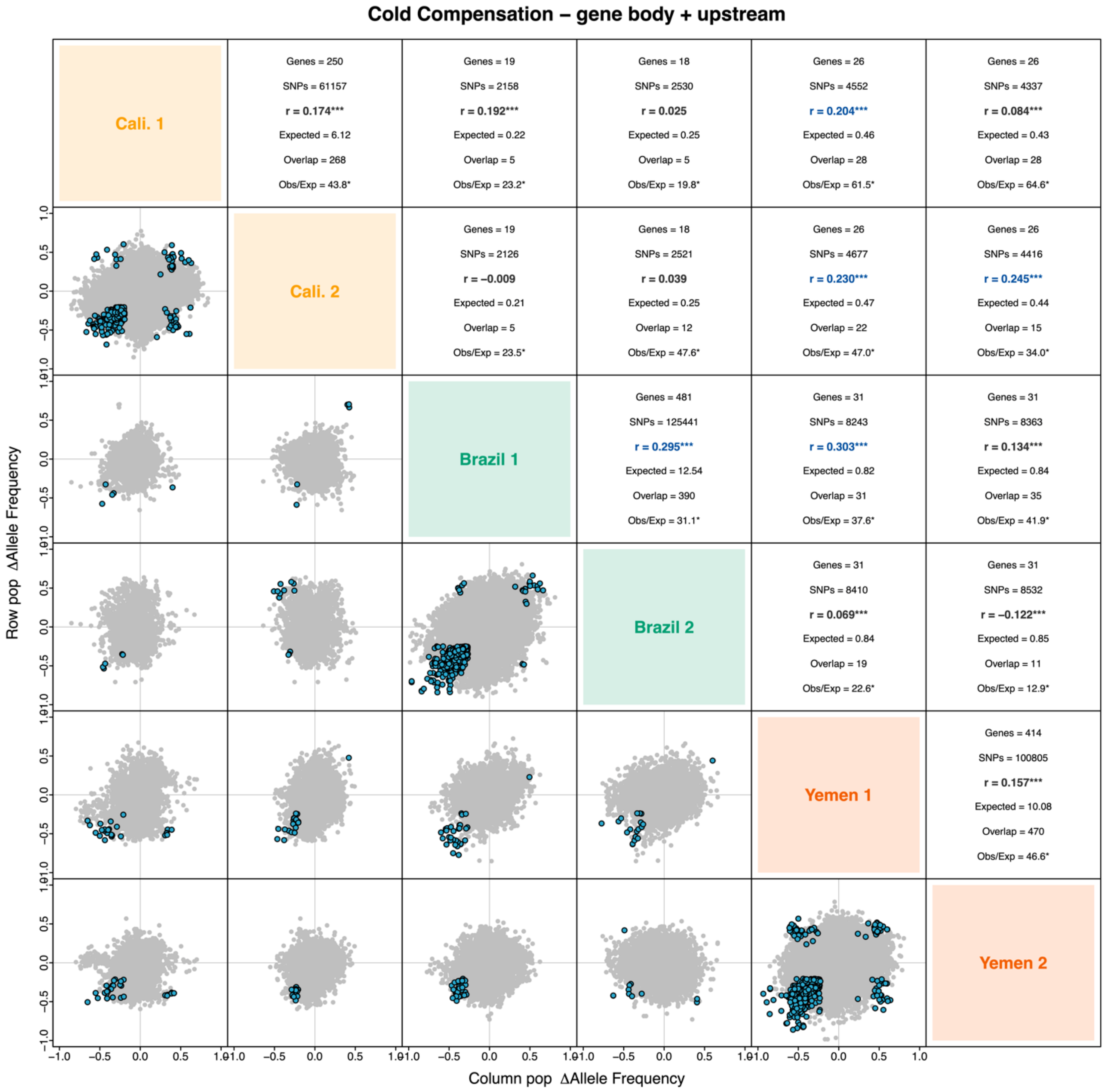


**Supplementary Figure 16: Repeatability of allele frequency change at cold compensation genes.** Shown are the observed and expected overlaps of outlier SNPs (top 1% in each population) between pairwise population replicates adapting to cold temperature. The compared SNPs are those that occurred in significantly differentially expressed genes detected on the backgrounds being compared (two backgrounds for between, and a single background for within comparisons). For cold compensation genes, the expected overlap was much greater than expected by chance for both within and between comparisons.

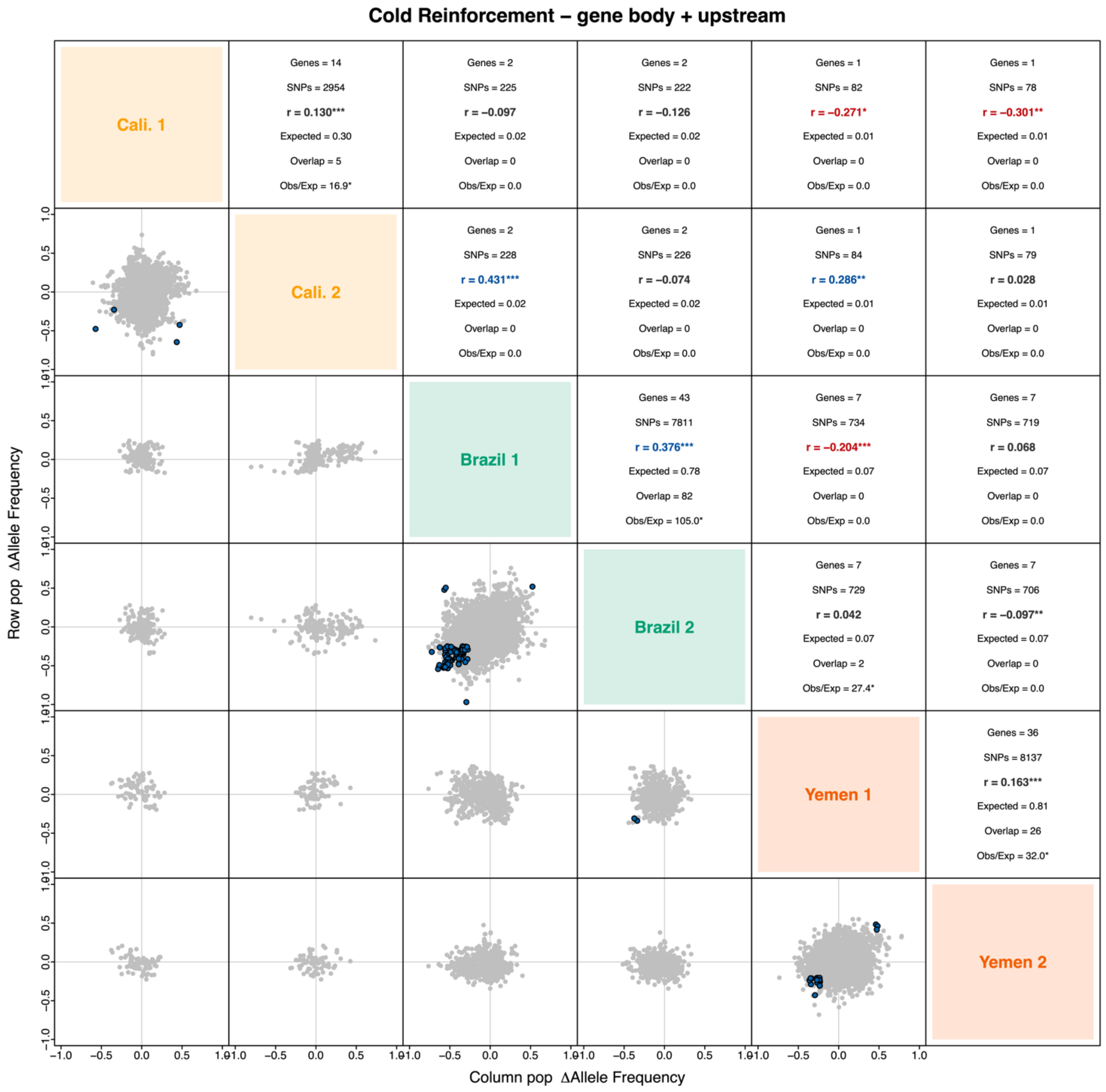


**Supplementary Figure 17: Repeatability of allele frequency change at cold reinforcement genes**. Shown are the observed and expected overlaps of outlier SNPs (top 1% in each population) between pairwise population replicates adapting to cold temperature. The compared SNPs are those that occurred in significantly differentially expressed genes detected on the backgrounds being compared (two backgrounds for between, and a single background for within comparisons). For cold reinforcement genes, the expected overlap was much greater than expected by chance for within comparisons, but largely non-existent for between comparisons, with very few SNPs entering the analysis overall.

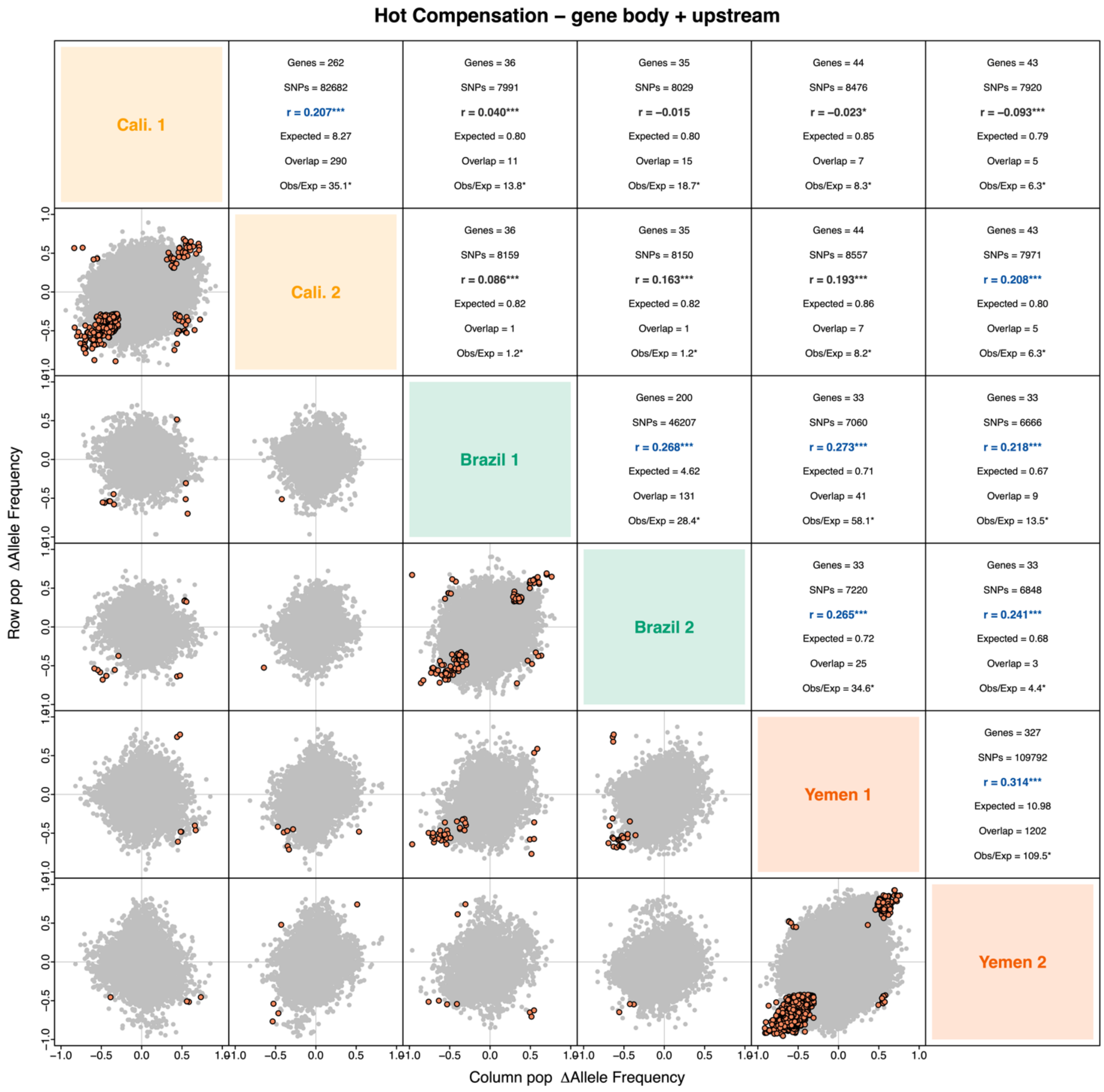


**Supplementary Figure 18: Repeatability of allele frequency change at heat compensation genes.** Shown are the observed and expected overlaps of outlier SNPs (top 1% in each population) between pairwise population replicates adapting to hot temperature. The compared SNPs are those that occurred in significantly differentially expressed genes detected on the backgrounds being compared (two backgrounds for between, and a single background for within comparisons). For heat compensation genes, the expected overlap was generally greater than expected by chance for both within and between comparisons, but much stronger for comparisons within background.

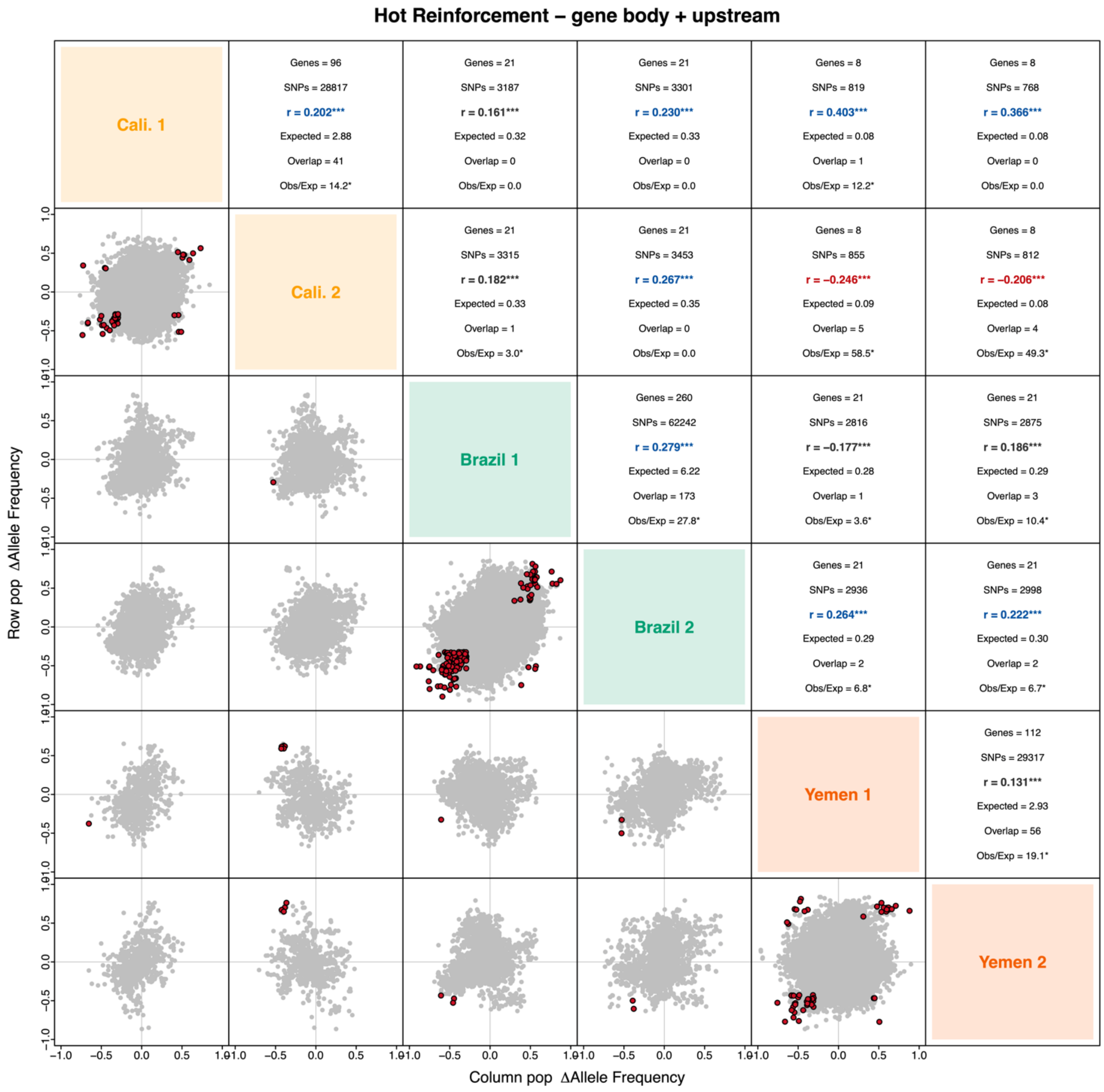


**Supplementary Figure 19: Repeatability of allele frequency change at heat reinforcement genes.** Shown are the observed and expected overlaps of outlier SNPs (top 1% in each population) between pairwise population replicates adapting to hot temperature. The compared SNPs are those that occurred in significantly differentially expressed genes detected on the backgrounds being compared (two backgrounds for between, and a single background for within comparisons). For heat reinforcement genes, the expected overlap was generally greater than expected by chance for both within and between comparisons, but stronger for comparisons within background.
